# A coevolution experiment reveals parallel mutations in the AcrA-AcrB-TolC efflux pump that contributes to bacterial antibiotic resistance

**DOI:** 10.1101/2022.03.16.484489

**Authors:** John L. Chodkowski, Ashley Shade

## Abstract

One interference mechanism of bacterial competition is the production of antibiotics. Bacteria exposed to antibiotics can resist antibiotic inhibition through intrinsic and/or acquired mechanisms. Here, we performed a coevolution experiment to understand long-term consequences of antibiotic production and antibiotic resistance for two environmental bacterial strains. We grew five independent lines of the antibiotic-producing environmental strain, *Burkholderia thailandensis* E264, and the antibiotic-inhibited environmental strain, *Flavobacterium johnsoniae* UW101, together and separately on agar plates for 7.5 months (1.5 month incubations), transferring each line five times to new agar plates. We first observed that the *F. johnsoniae* ancestor could tolerate the *B. thailandensis*-produced antibiotic through efflux mechanisms. We then sequenced the genomes of clonal isolates from the coevolved and monoculture *F. johnsoniae* lines, and uncovered mutational ramifications to the long-term antibiotic exposure. The coevolved genomes from *F. johnsoniae* revealed four potential mutational signatures of antibiotic resistance that were not observed in the evolved monoculture lines. Two mutations were found in *tolC*: one corresponding to a 33 bp deletion and the other corresponding to a nonsynonymous mutation. A third mutation was observed as a 1 bp insertion coding for a RagB/SusD nutrient uptake protein. The last mutation was a G83R nonsynonymous mutation in acetyl-coA carboxylayse carboxyltransferase subunit alpha (AccA). Placing the *tolC* 33 bp deletion back into the *F. johnsoniae* ancestor conferred some antibiotic resistance, but not to the degree of resistance observed in coevolved lines. Furthermore, the *accA* mutation matched a previously described mutation conferring resistance to *B. thailandensis*-produced thailandamide. Analysis of *B. thailandensis* transposon mutants for thailandamide production revealed that thailandamide was bioactive against *F. johnsoniae*, but also suggested that additional *B. thailandensis*-produced antibiotics were involved in the inhibition of *F. johnsoniae*. This study reveals how long-term interspecies chemical interactions can result in a novel mutation in efflux that contribute to antibiotic resistance.

## Introduction

Cultivation-independent sequencing of environmental microbial DNA has revealed the prevalence of antibiotic resistance genes in pristine environments (Allen at al. 2010), indicating that antibiotics and their corresponding resistance mechanisms have long-evolved in natural environments that predate their use in medicine (D’Costa et al. 2006). For example, glycopeptide antibiotics and resistance mechanisms have been present in bacterial genomes for at least 150 million years (Waglechner et al. 2019). Thus, it is expected that understanding the evolution of antibiotic resistance in “natural” settings will provide insights into emerging mechanisms of antibiotic resistance that may be useful for addressing resistance in clinical settings (Walsh 2013).

Microbial antibiotic production in the environment is typically viewed through the lens of competition (van der Meij et al. 2017). Bacteria produce antibiotics that interfere directly with competitors by inflicting cell damage (Hibbing et al. 2010). The DNA blueprints for antibiotics are organized in biosynthetic gene clusters (BSGC), in which locally proximal genes collectively encode the pathway for molecule production (Medema et al. 2015). The activation of BSGCs is typically tied to stress regulation (Baral et al. 2018), suggesting antibiotic production can be deployed as a survival strategy when conditions are not optimal for growth (Granato et al. 2019).

Bacteria can survive antibiotic exposure through the upregulation of intrinsic resistance mechanisms and can achieve antibiotic resistance through acquired mechanisms (e.g. Mutation or horizontal gene transfer; Arzanlou et al. 2017). Multidrug efflux pumps are particularly interesting because they can provide both intrinsic and acquired resistance mechanisms (Poole 2004). For example, low-levels of antibiotic exposure can upregulate intrinsic mechanisms of resistance, such as efflux pumps (Du et al. 2014; Frimodt-Møller and Løbner-Olesen 2019). The survival of bacteria to low levels of antibiotics can facilitate adaptive resistance (Ebbensgaard et al. 2020). This has been demonstrated by clinically relevant antibiotic resistance achieved via efflux pumps (Blair et al. 2014). Therefore, studying multigenerational interactions between antibiotic-producing and antibiotic-inhibited environmental isolates may provide insight into evolutionary dynamics driving antibiotic resistance.

We performed an agar-based experimental coevolution with two environmental strains, *B. thailandensis* and *F. johnsoniae*. These strains were co-plated together and, in parallel, also plated in monocultures on M9 minimal medium agar plates containing 0.2% glucose for over the span of 7.5 months. *B. thailandensis* and *F. johnsoniae* were co-plated such that *B. thailandensis* antibiotic inhibition of *F. johnsoniae* could occur without intergrowth of the colonies. By comparing outcomes of the coevolved lines to the evolved monoculture lines, we asked: What are the genetic and phenotypic repercussions of coevolution and how consistent are they across independent, replicate lines? What are the genetic signatures of *F. johnsoniae* antibiotic resistance? What is the antibiotic produced by *B. thailandensis* that inhibits *F. johnsoniae*? We found that coevolved *F. johnsoniae* lines gained resistance to the *B. thailandensis*-produced antibiotic while evolved monoculture lines remained susceptible. A 33 bp deletion in *tolC* and a nonsynonymous mutation in *accA* suggested two different paths to the evolution of this antibiotic resistance in *F. johnsoniae*. The ancestor *F. johnsoniae* strain became resistant after we deleted the *tolC* 33 bp region, but not to an equivalent level of resistance observed in the coevolved lines. This result suggests that multiple mutations may contribute to the observed resistance. Lastly, we provide evidence that thailandamide, a previously described antibiotic that inhibits fatty acid synthesis (Wu and Seyedsayamdost 2018), was among the bioactive compounds that inhibited *F. johnsoniae*. Our data also suggest that multiple antibiotics produced by *B. thailandensis* contributed to the observed inhibition of *F. johnsoniae*.

## Results

### *B. thailandensis* produces an antibiotic(s) that inhibits *F. johnsoniae*

We first observed that *B. thailandensis* inhibited *F. johnsoniae* when co-plated on M9-agar plates (Fig. 1). *B. thailandensis* exhibited radial growth on all edges along the circumference of the colony, while the *F. johnsoniae* colony proximal to *B. thailandensis* was inhibited. However, the distal end of the *F. johnsoniae* colony grew away from *B. thailandensis*, suggesting that *F. johnsoniae* may have intrinsic antibiotic resistance.

**Fig. 1.**
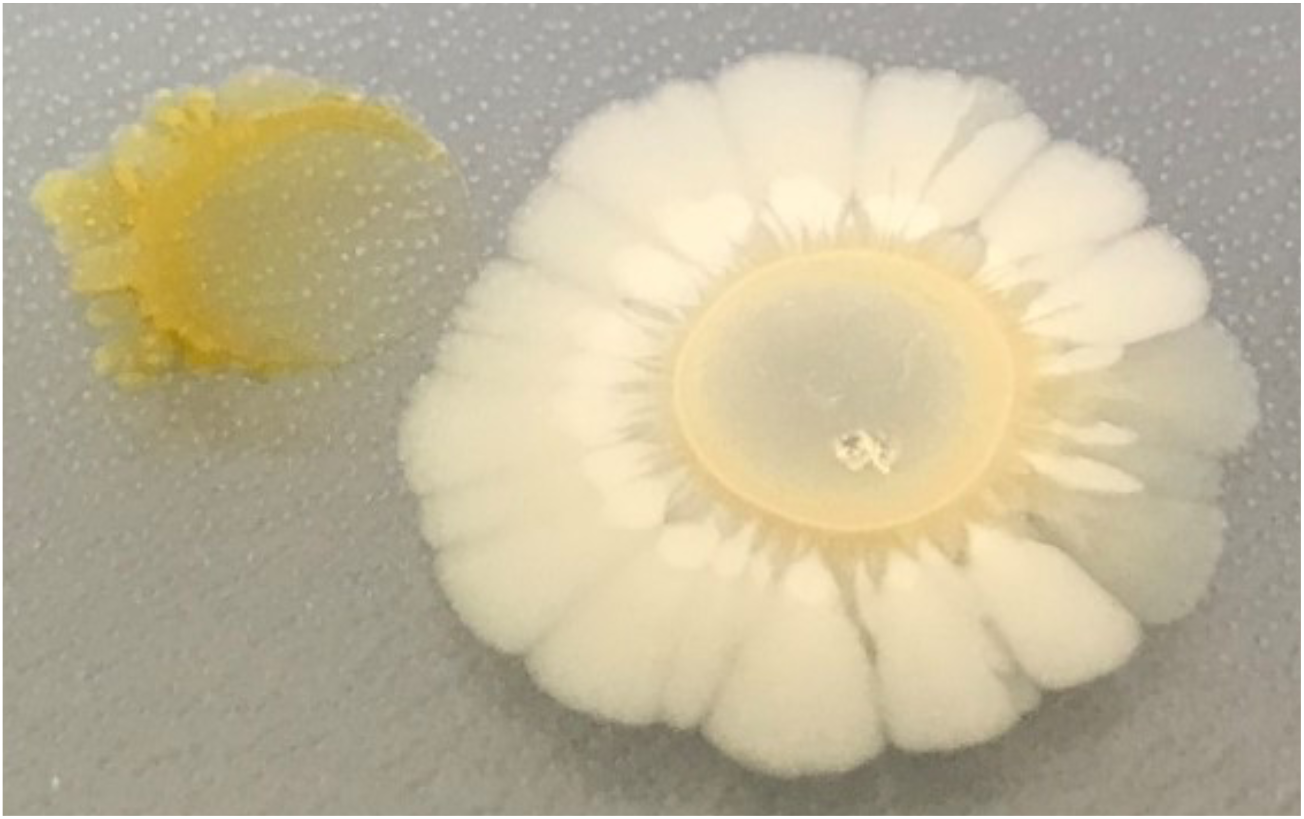
*B. thailandensis-*produced antibiotic(s) inhibits *F. johnsoniae. B. thailandensis* (right) and *F. johnsoniae* (left) were co-plated on M9-glucose agar at a distance that allowed for chemical interactions. An unidentified antibiotic(s) inhibited *F. johnsoniae*.

### Efflux allows *F. johnsoniae* to grow in the presence of a *B. thailandensis*-produced antibiotic

Given the growth pattern of *F. johnsoniae* when co-plated with *B. thailandensis*, we hypothesized that efflux contributed to *F. johnsoniae* intrinsic antibiotic resistance. We collected the organic fraction from spent supernatant of *B. thailandensis* grown in monoculture, which contained the antibiotic(s). We then treated *F. johnsoniae* cultures with the supernatant alone and in combination with daidzein, an efflux pump inhibitor (Aparna et al. 2014). While the supernatant inhibited *F. johnsoniae*, daidzein did not, and the supernatant with daidzein synergistically inhibited *F. johnsoniae* (Fig. 2, Supplementary Fig. 1). This suggested that *F. johnsoniae* had intrinsic antibiotic resistance via efflux-mediated antibiotic extrusion.

**Fig. 2.**
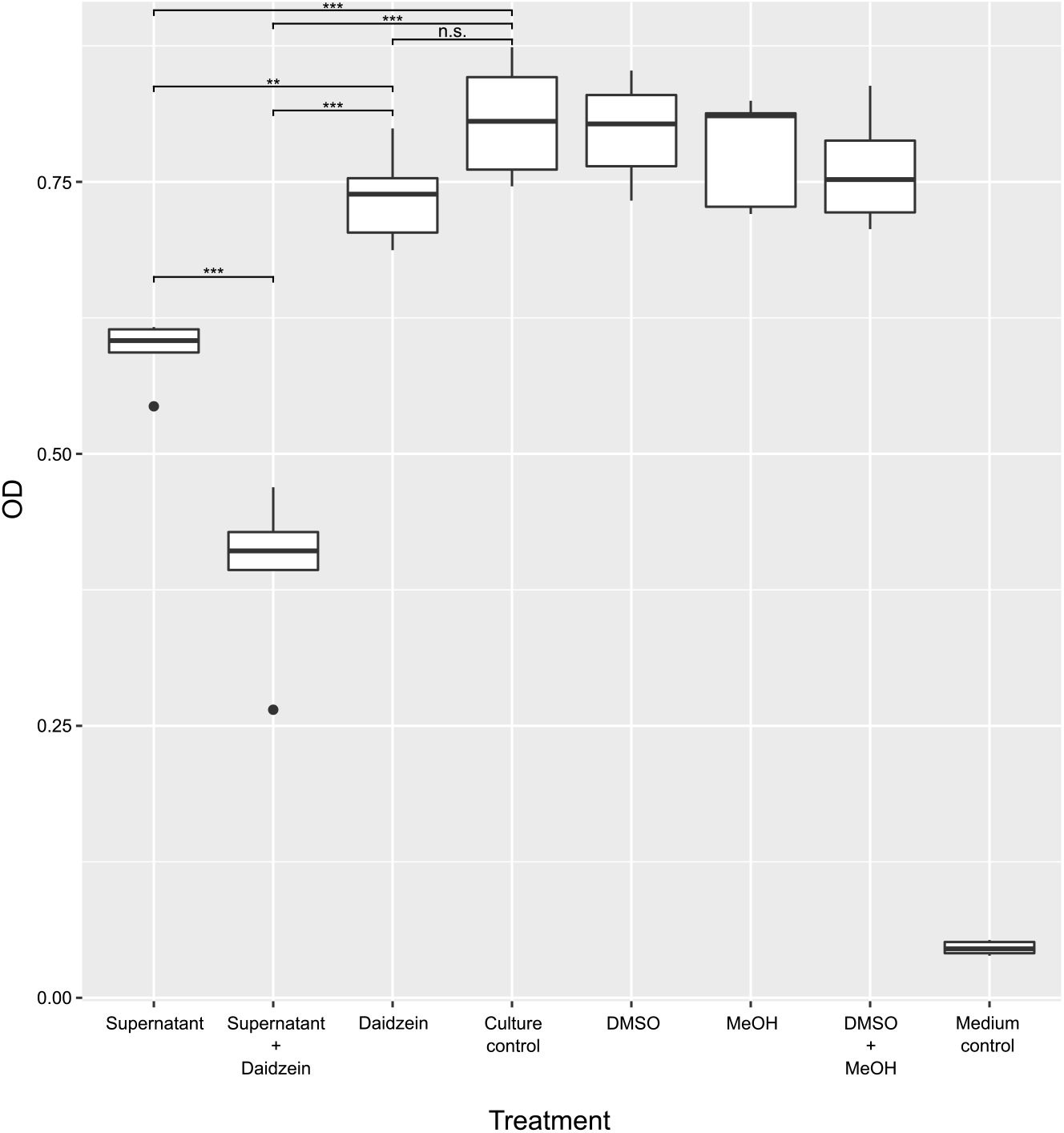
*F. johnsoniae* efflux system contributes to the extrusion of a *B. thailandensis*-produced antibiotic(s). An end-point growth measurement was taken after *F. johnsoniae* incubation with *B. thailandensis* culture supernatant, with the efflux pump inhibitor (daidzein), or with a combination of the supernatant and daidzein. An untreated *F. johnsoniae* culture, a *F. johnsoniae* culture with dimethyl sulfoxide (DMSO), a *F. johnsoniae* culture with methanol (MeOH), a *F. johnsoniae* culture with combined solvents, and blank medium served as controls. An ANOVA was performed comparing all treatments to the culture control. A Tukey HSD post-hoc analysis was performed for pairwise comparisons. Data shown are representative of 5 independent experiments; * p<0.05, ** p<0.001, ***p<0.0001, n.s.: not significant.

Because the supernatant was derived from *B. thailandensis* monoculture, we reasoned that production of the antibiotic(s) was not induced by the presence of *F. johnsoniae*, and instead, was likely constitutively produced or induced by abiotic factors or intrapopulation interactions. Bactobolin is well-characterized antibiotic that is produced by *B. thailandensis* in stationary phase monocultures (Duerkop et al. 2009; Amunts et al. 2015). Thus, we hypothesized that bactobolin was the antibiotic that inhibited *F. johnsoniae*. To test this hypothesis, we co-plated *F. johnsoniae* with a *B. thailandensis btaK*::T23 transposon mutant. Because *btaK* (BTH_II1233) is directly involved in the biosynthesis of bactobolin, *btaK* mutants do not produce bactobolin (Duerkop et al. 2009). However, we found that the *B. thailandensis btaK*::T23 transposon mutant inhibited *F. johnsoniae* despite the inability to produce bactobolin (Supplementary Fig. 2), suggesting either that bactobolin was ineffective against *F. johnsoniae* or that multiple antibiotics were involved in *F. johnsoniae* inhibition.

### Coevolutionary outcomes of *B. thailandensis*-*F. johnsoniae* interactions

To better understand the underlying factors contributing to *F. johnsoniae* resistance, we next asked how the antibiotic resistance of *F. johnsoniae* would change when coevolved with *B. thailandensis*. We performed an agar-based coevolution experiment with 10 µLvolume of two liquid overnight cultures spotted 14 mm apart to allow for extracellular chemical interactions, allowed to grow into colonies, and passaged onto another plate before intergrowth of the colonies could occur (Fig 3). The colonies spotted were either one of each of *B. thailandensis* and *F. johnsoniae* (“coevolved”), two of the same *B. thailandensis* colonies, or two of the same *F. johnsoniae* colonies (“monoculture”). All monoculture controls were grown in parallel to the co-evolved lines (Fig. 3). We observed that, while *B. thailandensis* substantially inhibited *F. johnsoniae* on the first plate (Supplementary Fig. 3), the radial growth of *F. johnsoniae* generally increased with each successive plate passage (Fig. 4, Supplementary Fig. 3), suggesting that co-evolved lines of *F. johnsoniae* were gaining antibiotic resistance. Indeed, when a freezer stock from a coevolved *F. johnsoniae* line and a freezer stock from the corresponding monoculture *F. johnsoniae* line were plated with the *B. thailandensis* ancestor, *F. johnsoniae* from the coevolved line displayed more radial growth and thus more resistance as compared to the evolved monoculture control (Supplementary Fig. 4). Interestingly, colonies from the coevolved *F. johnsoniae* lines were also able to resist colony invasion by *B. thailandensis*, while colonies from the evolved monoculture lines could not (Supplementary Fig. 5). Overall, these results suggest that increased resistance to antibiotics in *F. johnsoniae* evolved as an outcome of long-term exposure to and engagement with *B. thailandensis*.

**Fig. 3.**
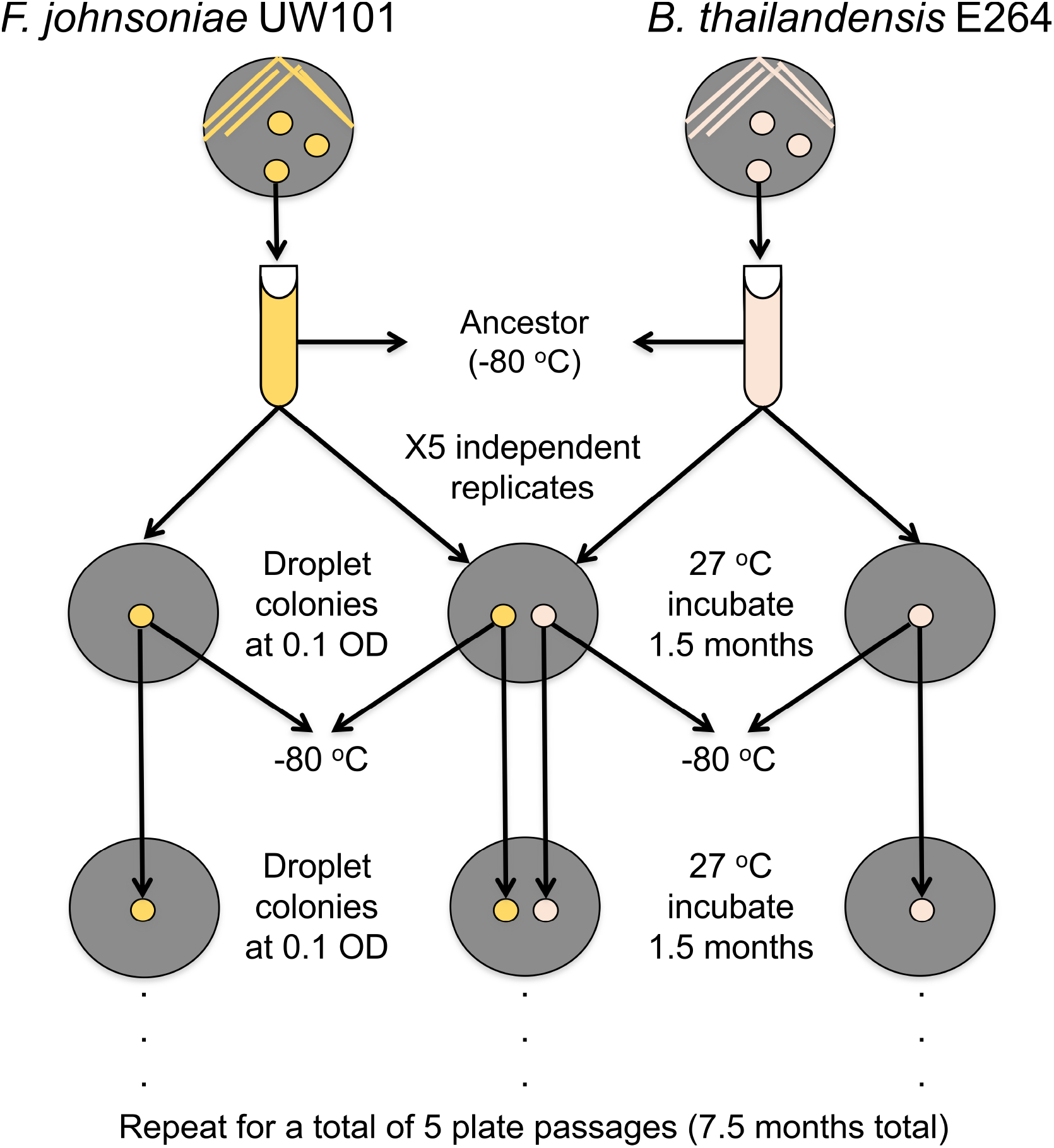
Schematic of (co)evolution experiment.

**Fig. 4.**
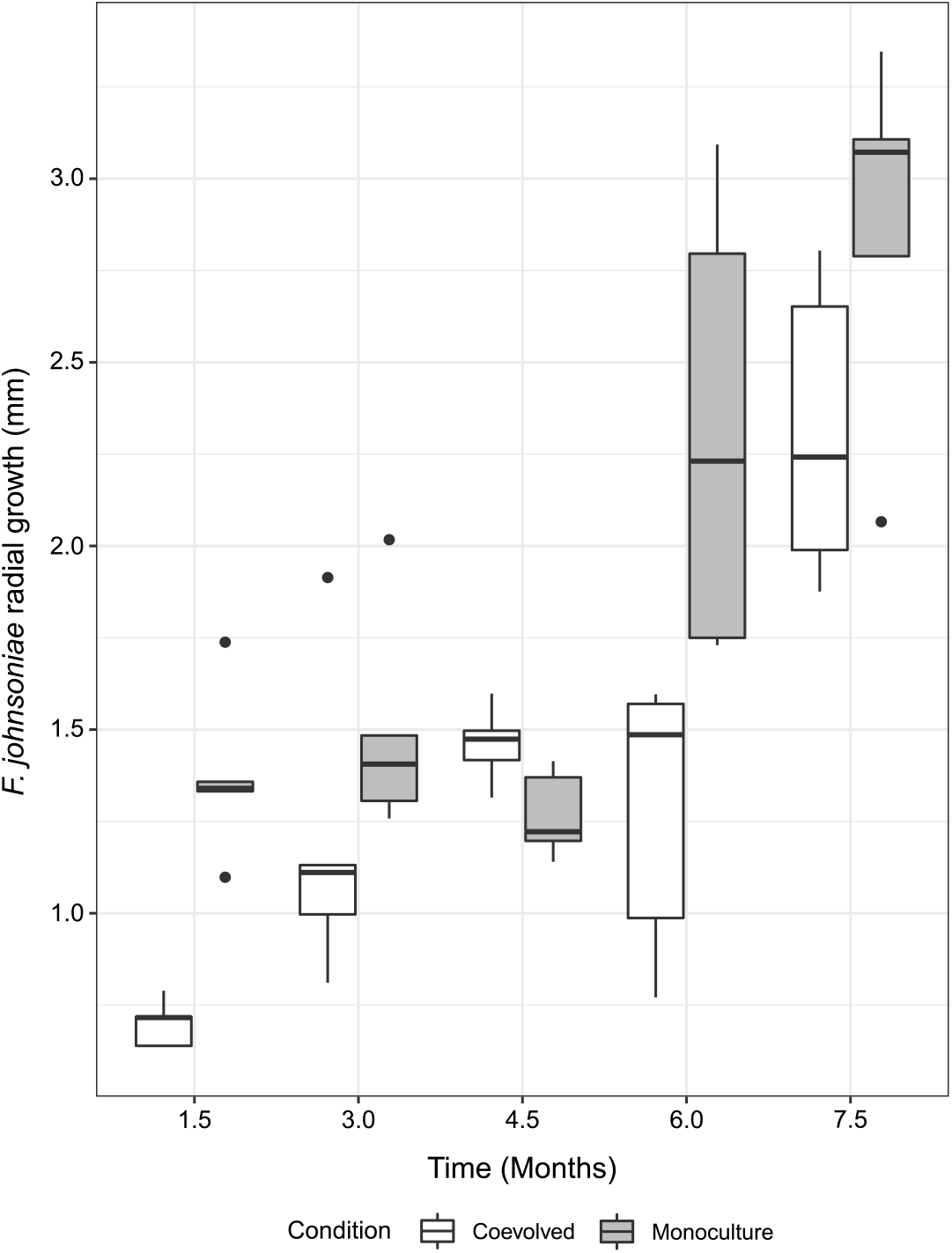
*F. johnsoniae* trends toward increased growth success with each plate passage in the presence of *B. thailandensis*. Radial colony growth of coevolved *F. johnsoniae* was measured from the center of the colony to point of furthest growth on the agar plate. Measurements were taken at the end of each plate passage.

### Whole genome sequencing reveals genetic signatures of antibiotic resistance

We performed whole genome sequencing to discover mutations that may have contributed to increased antibiotic resistance in the *F. johnsoniae* coevolved lines. We sequenced clonal isolates from the *F. johnsoniae* ancestor and from 5 independent, replicate lines of coevolved and monocultures from the final (fifth) plates. We detected mutations in all genomes as compared to the ancestor (Table 1, Supplemental File 1). *F. johnsoniae* lines coevolved with *B. thailandensis* acquired mutations that were distinct from acquired mutations in *F. johnsoniae* lines grown in monocultures (Supplementary Fig. 6).

**Table 1.**
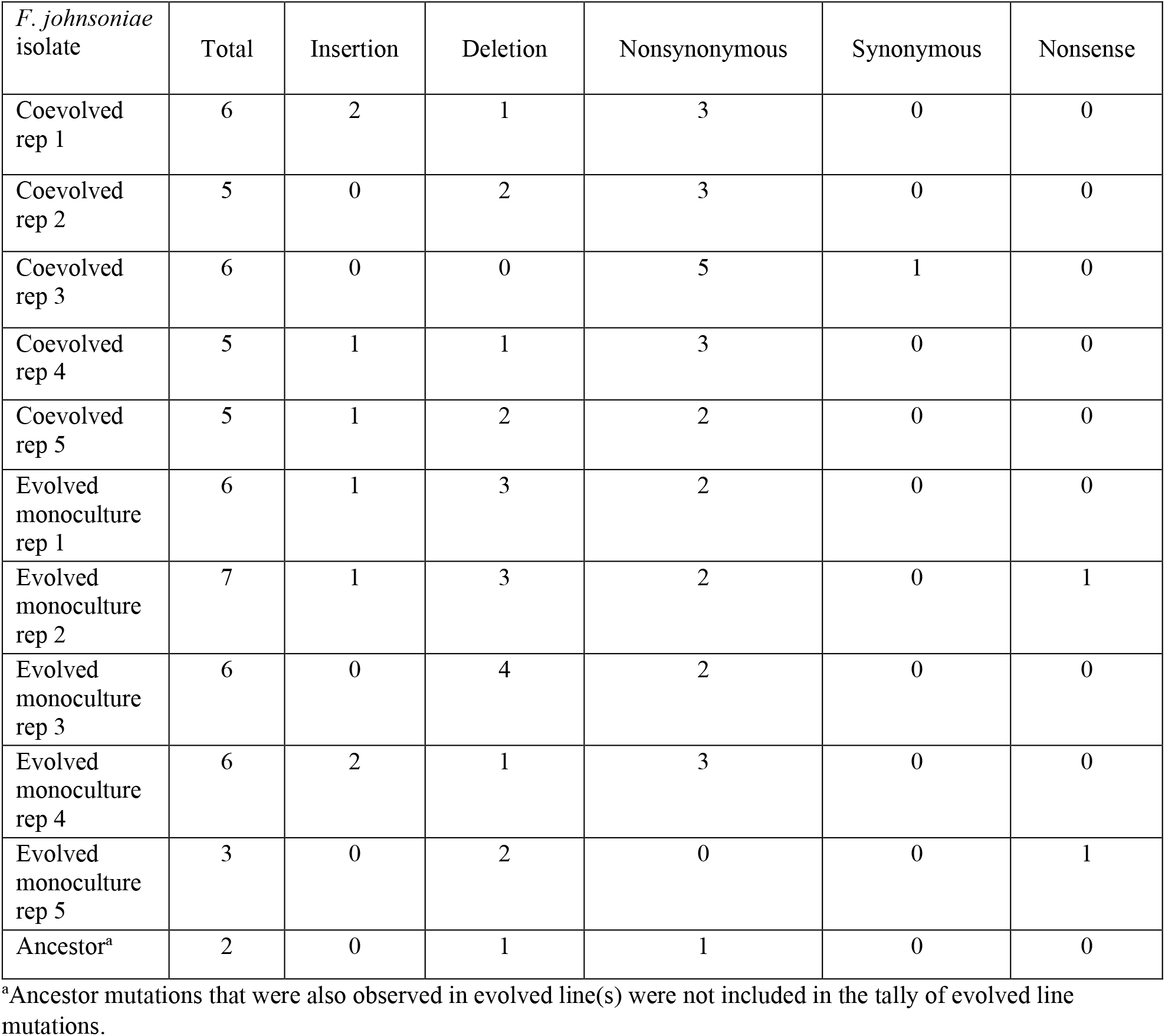
Summary of mutation types observed in *F. johnsoniae* from the (co)evolution experiment. Mutations are from five representative clonal isolates after the final (5^th^) plate passage and the ancestor.

Mutations in an efflux outer membrane protein again suggested a mechanism of acquired antibiotic resistance (Table 2). A *tolC* mutation was observed in 4 out of the 5 coevolved lines. While *F. johnsoniae* contains 16 *tolC* genes (Supplementary Table 1), the same gene was mutated in 4 independent replicates. Furthermore, there was evidence of parallel evolution at the nucleotide level, as 3/4 coevolved lines with mutations in *tolC* had the same 33 bp deletion (Supplemental File 1). This deletion was from nucleotides 261-293 in the FJOH_RS06580 coding sequence, resulting in an in-frame deletion (Fig. 5). This deletion affected one of the extracellular loops of TolC. The WT sequence revealed a 11 bp direct repeat, occurring before the deletion and representing the last 11 bp of the 33 bp deletion. This suggests that the deletion may have occurred during a replication deletion event. The other mutation in FJOH_RS06580 (coevolved line 3) was a single nucleotide polymorphism (G247A in the coding sequence) that resulted in a nonsynonymous mutation (G83R). Protein modeling suggested that this mutation resulted in a decreased diameter of the efflux channel (Supplementary Fig. 7). Coevolved line 4 did not harbor a mutation in FJOH_RS06580 but instead had a unique bp insertion in a *ragB*/*susD* nutrient uptake outer membrane protein (FJOH_RS24865). Guanine was inserted at nucleotide position 794 in the coding sequence for FJOH_RS24865. This frameshift mutation resulted in a premature termination codon 6 bp downstream of the insertion, likely rendering the protein nonfunctional.

**Table 2.** Distinctions and overlaps of loci with mutations unique to the *F. johnsoniae* lines that were coevolved with *B. thailandensis*. These mutations were detected (+) in at least one coevolved line and not present (-) in any of the evolved *F. johnsoniae* monocultures

| *F. johnsoniae*<br>Locus | Annotation | Rep<br>1 | Rep<br>2 | Rep<br>3 | Rep<br>4 | Rep<br>5 |
| --- | --- | --- | --- | --- | --- | --- |
| FJOH_RS00255 | acetyl-CoA carboxylase carboxyltransferase subunit alpha | - | - | + | - | - |
| FJOH_RS02320 | aminomethyl-transferring glycine dehydrogenase | - | - | - | - | + |
| FJOH_RS04780 | hypothetical protein | + | - | - | - | - |
| FJOH_RS06580 | TolC family protein | + | + | + | - | + |
| FJOH_RS07830 | NAD(P)/FAD-dependent oxidoreductase | - | + | - | - | - |
| FJOH_RS09515 | response regulator transcription factor | - | + | - | - | + |
| FJOH_RS09520 | GHKL domain-containing protein | + | - | + | + | - |
| FJOH_RS11170 | Gfo/Idh/MocA family oxidoreductase | - | - | + | - | - |
| FJOH_RS12240 | phosphoribosylformylglycinamide cyclo-ligase | - | - | + | - | - |
| FJOH_RS14175 | response regulator transcription factor | - | + | - | - | - |
| FJOH_RS20510 | TetR/AcrR family transcriptional regulator | + | - | - | - | - |
| FJOH_RS21875 | glycoside hydrolase | + | - | - | - | - |
| FJOH_RS24865 | RagB/SusD family nutrient uptake outer membrane protein | - | - | - | + | - |
| FJOH_RS25290 | phosphoribosylanthranilate isomerase | - | - | - | + | - |

**Fig. 5.**
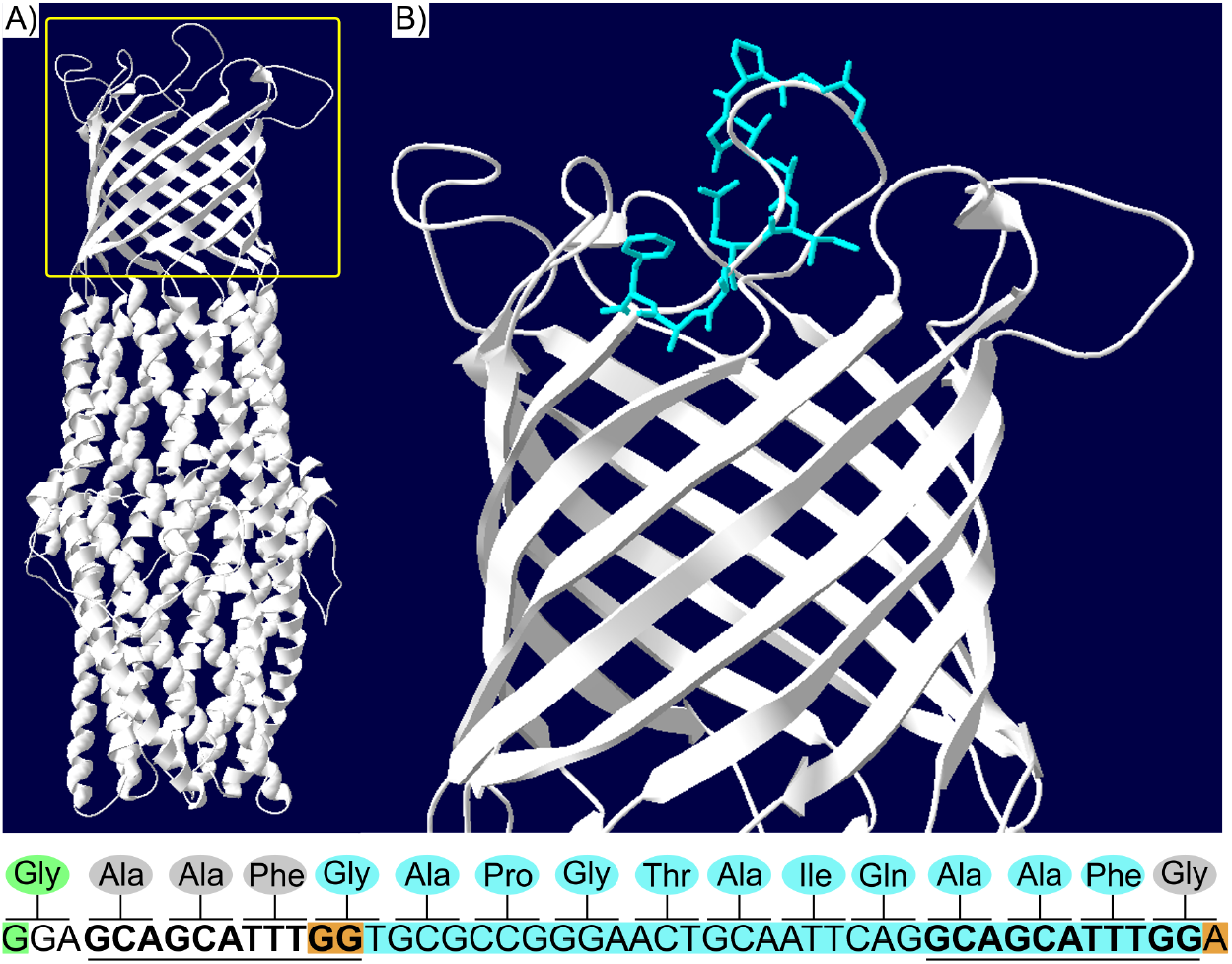
A *tolC* 33 bp deletion is located on a TolC extracellular loop. The TolC protein (A) contains a α-helical trans-periplasmic tunnel, a β-barrel channel embedded in the outer membrane, and extracellular loops at the cell surface. The yellow box represents the inset in (B), where the 11 amino acids corresponding to the 33 bp deletion are highlighted in blue. These amino acids are part of one of the extracellular loops. The nucleotide sequence is shown below the image, representing bps 247-294 in the FJOH_RS06580 coding sequence. The corresponding amino acids are shown above each codon. Nucleotides highlighted in blue represent the 33 bp deletion. Nucleotides highlighted in orange show how the deletion was in-frame. The 11 bp direct repeats are in bold and underlined. The nucleotide highlighted in green is associated with the nonsynonymous mutation (G247A in the coding sequence, G83R in TolC)

### The FJOH_RS06580 *tolC* 33 bp deletion confers antibiotic resistance

We asked whether the mutations in *tolC* or the mutation in *ragB/susD* of *F. johnsoniae* would provide resistance to the *B. thailandensis*-produced antibiotic(s). These mutations were amplified from the coevolved *F. johnsoniae* lines and recombined into the ancestor so that the ancestor would only harbor one of these mutations and not any of the additional mutations observed in the coevolved lines. Strains and plasmids used to make recombinant *F. johnsoniae* are outlined in Table 3. Ancestor *B. thailandensis* inhibited the *F. johnsoniae* ancestor and recombinant strains JCAC03 and JCAC04 (Fig. 6). In fact, JCAC03 appeared more inhibited compared to the *F. johnsoniae* ancestor, suggesting that a decreased diameter to the TolC efflux channel (inferred from protein structure predicted by SWISS-MODEL) is detrimental to fitness. In contrast, recombinant strain JCAC01 (*tolC* 33 bp deletion) was the least inhibited when plated with ancestor *B. thailandensis*, suggesting that the 33 bp deletion confers some degree of antibiotic resistance (Fig. 6B). Thus, it appears that *F. johnsoniae* uses efflux pumps for intrinsic resistance and three of the coevolved clonal isolates from *F. johnsoniae* acquired a mutation in an efflux pump that conferred increased antibiotic resistance to a *B. thailandensis*-produced antibiotic(s). However, the 33 bp deletion in *tolC* alone did not confer equivalent resistance observed in the coevolved lines, as recombinant strain JCAC01 was resistant in the presence of *B. thailandensis*, but not as resistant as a coevolved line harboring the *tolC* 33 bp deletion with additional mutations (Supplementary Fig. 8). This suggests that coevolved lines harboring the *tolC* 33 bp mutation may contain synergistic antibiotic resistance-conferring mutations.

**Table 3.**
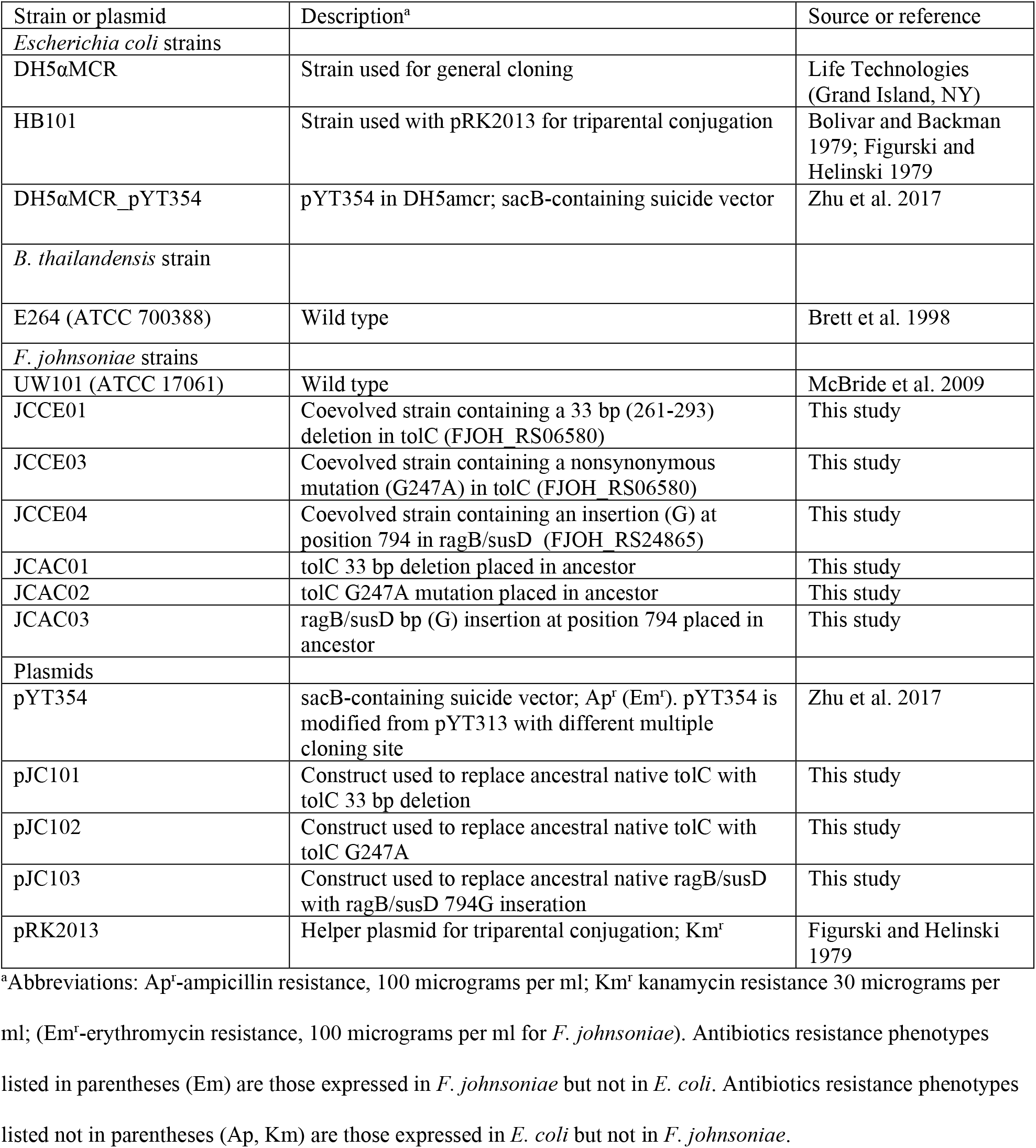
Strains and plasmids used in this study.

**Fig. 6.**
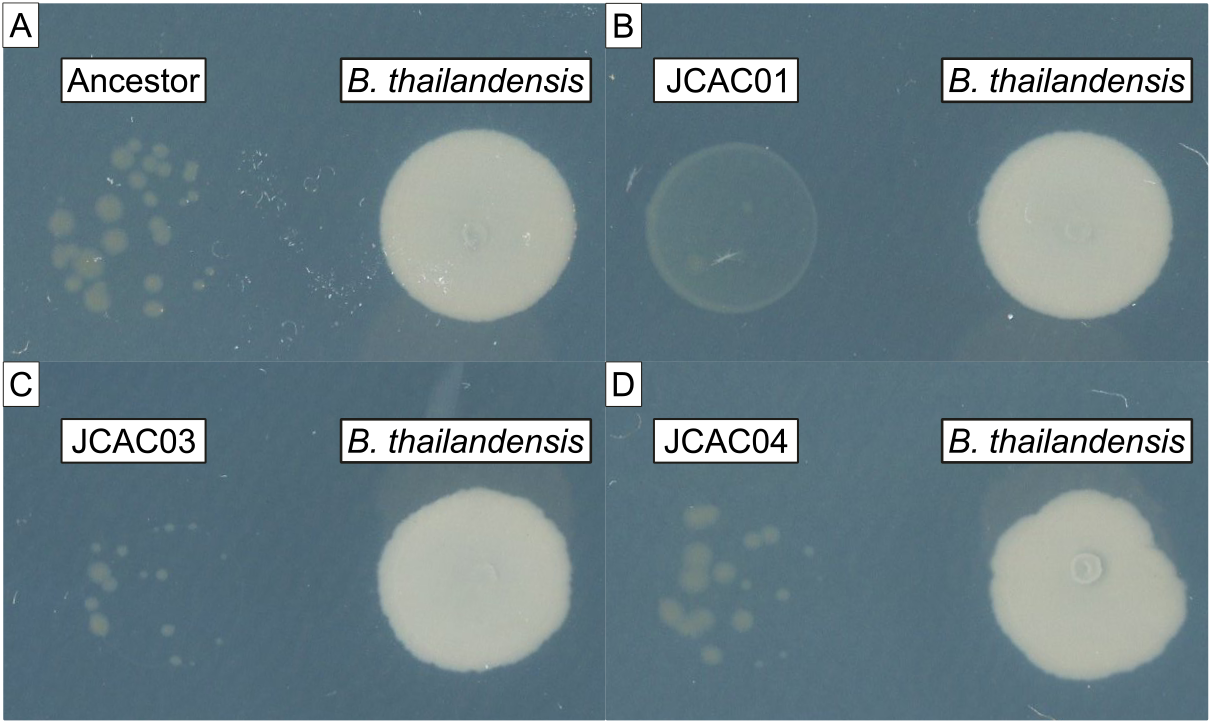
The *tolC* 33 bp deletion confers antibiotic resistance. The ancestor (A) and recombinant strains of *F. johnsoniae* ancestor (C-D) were co-plated with *B. thailandensis*. The recombinant strain with the *tolC* 33 bp deletion is the least inhibited by *B. thailandensis* compared to the ancestor and other recombinant strains. Plates were imaged after a week of incubation.

### Thailandamide is one of the bioactive molecules that inhibits *F. johnsoniae*

The nonsynonymous mutation in *F. johnsoniae* coevolved line 3 *tolC* did not confer resistance to the *B. thailandensis*-produced antibiotic(s). But, *F. johnsoniae* coevolved line 3 still developed resistance to the antibiotic(s) despite harboring this mutation (Fig. 7A). This *F. johnsoniae* coevolved line 3 also contained a unique nonsynonymous mutation in acetyl-CoA carboxylase carboxyltransferase subunit alpha (*accA*, Table 2). This was a C479A nonsynonymous mutation in the coding sequence of *accA* that resulted in a P160Q alteration in AccA. This mutation was similar to a P164Q mutation in AccA from *Salmonella enterica* serovar Typhimurium strain LT2 that conferred resistance to thailandamide from *B. thailandensis* (Wozniak et al. 2018). The P164Q mutation from *S. enterica* aligned with the P160Q mutation observed in our coevolved line (Fig. 7). Thus, we hypothesized that thailandamide was one of the antibiotics responsible for inhibition of *F. johnsoniae*.

**Fig. 7.**
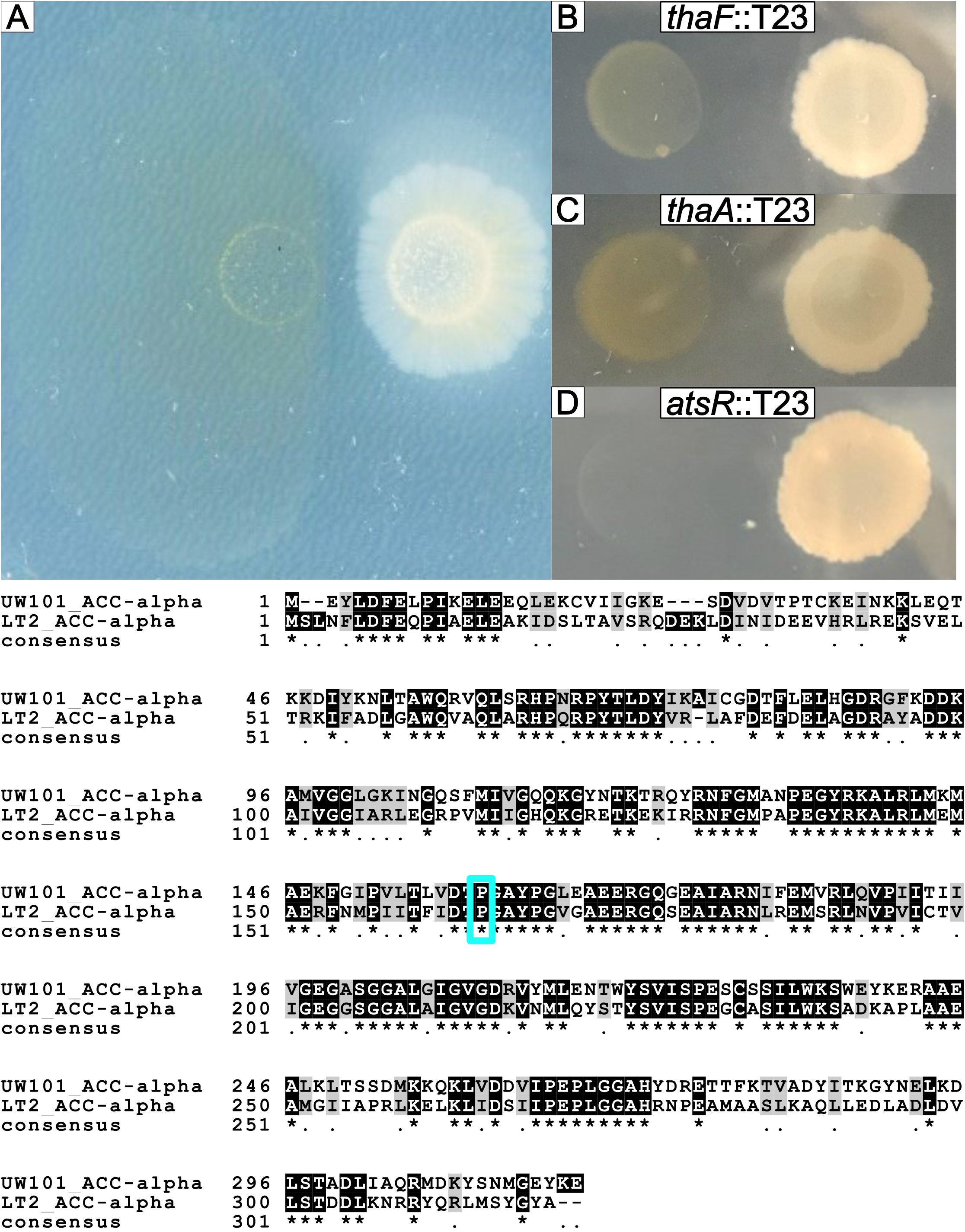
Thailandamide is bioactive against *F. johnsoniae*. While the nonsynonymous mutation in *tolC* did not confer antibiotic resistance (present in *F. johnsoniae* coevolved line 3), *F. johnsoniae* coevolved line 3 still displayed antibiotic resistance at the end of plate passage five (A). *B. thailandensis* transposon mutants in *thaF* (B) and *thaA* have decreased inhibition toward *F. johnsoniae* while an *atsR* transposon mutant has increased inhibition of *F. johnsoniae* (D). Below the panels is an amino acid sequence alignment between the *F. johnsoniae* AccA and S. enterica AccA. The asterisks indicate positions within the proteins that were identical. The blue box highlights the alignment of P160 in *F. johnsoniae* and P164 in *S. enterica*.

We plated the *F. johnsoniae* ancestor with *B. thailandensis thaF* (BTH_II1675), *thaA* (BTH_II1681), and *atsR* (BTH_I0633) transposon mutants. *thaF* encodes a polyketide synthetase trans-AT domain directly involved in the biosynthesis of thailandamide, *thaA* encodes a LuxR-type regulator in the thailandamide biosynthetic gene cluster, and *astR* encodes a global regulator. ThaA positively regulates the thailandamide biosynthetic gene cluster while AtsR negatively regulates the thailandamide biosynthetic gene cluster. We found that *thaF*::T23 and *thaA*::T23 mutants decreased inhibition of *F. johnsoniae* (Fig. 7B, Fig. 7C) while the *atsR*::T23 mutant increased inhibition of *F. johnsoniae* (Fig. 7D). This suggests that thailandamide is bioactive against *F. johnsoniae*. However, *F. johnsoniae* was still slightly inhibited when plated with *thaF*::T23 and *thaA*::T23 mutants, which also suggests that in addition to thailandamide, there may be another *B. thailandensis*-produced antibiotic that is inhibiting *F. johnsoniae*. However, we were unable to determine this molecule.

## Discussion

We performed an experimental coevolution study between a strain capable of antibiotic production (*B. thailandensis*) and an antibiotic-inhibited strain (*F. johnsoniae*). Our findings show how parallel evolution, both at the gene level and in mutations, for antibiotic resistance occurred from long-term interspecies interactions facilitated through chemical exchange via diffusion in agar. In addition, we demonstrated that efflux provided both intrinsic and acquired mechanisms of antibiotic resistance. Analysis of mutations of *F. johnsoniae* coevolved lines as compared to lines evolved in monoculture revealed that antibiotic resistance was conferred via efflux. Specifically, a 33 bp deletion in *tolC*, which eliminated 11 amino acids that were part of an extracellular loop in TolC, conferred some, but not complete, resistance in the coevolved *F. johnsoniae*. A different path to resistance was achieved through a nonsynonymous mutation in *accA*. Furthermore, this *accA* mutation provided evidence that one of the *B. thailandensis*-produced antibiotics was thailandamide, given that the same mutation was described in a *Salmonella* strain that also gained resistance to thailandamide (Wozniak et al. 2018). *B. thailandensis* transposon mutants with abrogated thailandamide production confirmed that thailandamide was bioactive against *F. johnsoniae*, but slight inhibition was observed, again suggesting that multiple antibiotics are inhibiting *F. johnsoniae*.

Performing the experimental coevolution on agar plates created a heterogenous environment that preempted the evolution of antibiotic resistance. *F. johnsoniae* growth at the distal end of the colony was permitted because of an antibiotic concentration gradient that was established via diffusion. Low-dose antibiotics likely upregulated intrinsic mechanisms of resistance (e.g. efflux pumps) that conferred low-levels of resistance (Sandoval-Motta and Aldana 2016). The ability to survive exposure to antibiotics via intrinsic mechanisms (Frimodt-Møller et al. 2018; Meouche & Dunlop 2018) can provide the opportunity for mutational acquisition of resistance (Frimodt-Møller and Løbner-Olesen 2019; Ebbensgaard et al. 2020). This was demonstrated experimentally in a seminal study that found that a heterogenous environment increased the rate of adaptation to antibiotics with as few at 100 bacteria in the initial inoculum (Zhang et al. 2011). This approach has been expanded to show how the initial adaptation to low levels of antibiotics facilitate adaptations to high levels of resistance (Baym et al. 2016). Thus, evolutionary adaptations to antibiotic resistance can be fostered in heterogenous environments that would otherwise not be achieved in a uniform environment (Hermsen and Hwa 2010; Hermsen et al. 2012).

*F. johnsoniae* antibiotic resistance was conferred through a 33 bp deletion in *tolC*. The occurrence of the 11 bp directed repeats may support a replication misalignment event that led to the 33 bp deletion (Kong and Masker 1994; Bzymek and Lovett 2001). TolC forms the outmembrane channel part of the tripartite AcrAB-TolC drug efflux pump (Du et al. 2018). *tolC* (FJOH_RS06580) is located in an operon that includes the remaining components necessary for a functional efflux pump. This includes a TetR/AcrR family transcriptional regulator (FJOH_RS06575), a membrane fusion protein (FJOH_RS06585), and an inner membrane transporter (FJOH_RS0690). Elimination of 11 amino acids from an extracellular loop in TolC may result in TolC more frequently adopting an open conformation, which could increase the rate of antibiotic extrusion. “Leaky” TolC mutants have been characterized, but these mutations occurred at the periplasmic end of TolC (Bavro et al. 2008; Pei et al. 2011). In fact, mutational studies of the TolC extracellular loops appear uncommon but may present a novel mechanism for antibiotic resistance (Krishnamoorthy et al. 2013).

The *F. johnsoniae* recombinant strain harboring the 33 bp deletion in *tolC* was resistant, but not as resistant as the *F. johnsoniae* coevolved lines to the *B. thailandensis*-produced antibiotic(s). Additional mutations in the coevolved lines are likely to provide increased antibiotic resistance. For example, coevolved line 1 also harbored a 1 bp insertion (T) at position 4734734 in FJOH_RS20510, annotated as a TetR/AcrR family transcriptional regulator. This results in a nonsense mutation that would render the protein nonfunctional. TetR regulators are typically negative regulators, so a nonfunctional TetR regulator would lead to increased expression of efflux pump systems (Cuthbertson and Nodwell 2013). We note that this occurred in FJOH_RS20510 and not FJOH_RS06575, but some TetR regulators have multiple targets and TetR from FJOH_RS20510 could also be negatively regulating the FJOH_RS06580-FJOH_RS06585-FJOH_RS06590 efflux system (Colclough et al. 2019). The remaining coevolved lines with the 33 bp deletion in *tolC* (line 2 and line 5) also had mutations in transcriptional regulators (OmpR and LytTR) but their potential contributions to increased antibiotic resistance is unclear.

A nonsynonymous mutation in *accA* in coevolved line 3 also provided resistance to the *B. thailandensis*-produced antibiotic(s). In addition, this mutation guided our efforts to uncover that thailandamide was one of the antibiotics produced in *B. thailandensis* that inhibited *F. johnsoniae*. Thailandamide resistance was first characterized in spontaneous mutants from *S. enterica* (Wozniak et al. 2018). Wozniak and colleagues found 6 unique mutations for 3 different amino acid positions in AccA that provided thailandamide resistance. One of these mutations was also spontaneously generated in our study (P160Q in *F. johnsoniae*, P164Q in *S. enterica*). *B. thailandensis* transposon mutants with abrogated thailandamide production effectively reduced inhibition of *F. johnsoniae*. But, we still observed inhibition with the thailandamide transposon mutants. It is likely that *F. johnsoniae* was subjected to multiple antibiotics but other mutations conferring antibiotic resistance in the coevolved lines did not provide insight into other *B. thailandensis*-produced antibiotics affecting *F. johnsoniae*.

*F. johnsoniae* coevolved line 4 did not harbor a mutation in *tolC* or *accA*. However, the *ragB/susD* mutation was unique to coevolved line 4 and we hypothesized that this may provide an alternative mechanism for antibiotic resistance. Some antibiotics enter bacterial cells via nutrient transporters (Yoneyama and Nakae 1993; Castañeda-García et al. 2009). Since the *ragB/susD*, rendered the protein nonfunctional, antibiotic resistance could have been conferred to coevolved line 4 by prevention of antibiotic uptake. We did not find this to be the case, as the *F. johnsoniae* recombinant strains with the *ragB/susD* nonsynonymous mutation had equivalent susceptibility to the *B. thailandensis*-produced antibiotic(s) as the *F. johnsoniae* ancestor. The other mutations in coevolved line 4 may confer antibiotic resistance or, it is possible that mutations for antibiotic resistance did not fixate in the population, and we chose an isolate for sequencing that did not acquire a mutation conferring antibiotic resistance. Bacterial population sequencing could shed light on this discrepancy.

The diversity of antibiotics (Rutledge and Challis 2015) and corresponding resistance mechanisms (Mungan et al. 2020) demonstrate the breadth of genetic novelties that have arisen from millions of years of bacterial competition. Our results provide a case study of bacterial interspecies chemical engagements that can evolve in a heterogeneous environment, with insights into understanding the complex and potentially interacting mutational paths to antibiotic resistance. We were not able to identify the other antibiotic(s) effective against *F. johnsoniae* and we were not able to directly confirm that thailandamide is bioactive against *F. johnsoniae*. Antibiotic isolation and screening can be performed in future studies to discover and confirm our findings. In addition, future studies can perform population sequencing across the longitudinal study, lending insight into the mutational trajectories of the evolving populations.

## Materials and Methods

### Extraction of *B. thailandensis* supernatant containing antibiotic activity

A freezer stock of *B. thailandensis* was plated on 50% trypticase soy agar (TSA50) and grown overnight at 27 °C. A collection of lawn growth was transferred to 7 mL of M9 minimal salts-glucose (M9-glucose) medium to achieve an initial OD of ∼0.2. The culture was incubated over night at 27 °C, 200 rpm. The next day, 1 mL of culture was transferred to 50 mL fresh M9-glucose medium. The culture was incubated for 24 h at 27 °C, 200 rpm. The culture was transferred to a falcon tube and centrifuged at 4 °C, 5000 rpm for 20 min. The supernatant was removed, filtered with a 0.22 µM filter, and transferred to a separatory funnel. Ten mL of dichloromethane (DCM) was added to the separatory funnel. The separatory funnel was agitated three times and the DCM layer was removed. This process was repeated twice more in batches of 10 mL additions of DCM. The collected DCM layer was dried under nitrogen gas. The dried DCM extracts were reconstituted in 1 mL of a 50:50 methanol:water mixture. This mixture was used for the efflux pump inhibitor experiment.

### Efflux pump inhibitor experiment

A freezer stock of *F. johnsoniae* was plated on TSA50 and grown over night at 27 °C. Five individual colonies were then inoculated in 5 mL 50% trypticase soy broth (TSB50) and grown overnight at 27 °C, 200 rpm. After ∼16 hr growth, 50 µL of culture was diluted into 4.95 mL of fresh TSB50 for each independent replicate. Then, 650 µL aliquots were dispensed in each of seven tubes for each independent replicate. Seven conditions were prepared across the seven tubes as follows: 1) *F. johnsoniae* culture control 2) *F. johnsoniae* with DMSO control 3) *F. johnsoniae* with 50:50 methanol:water control 4) *F. johnsoniae* with DMSO + 50:50 methanol:water control 5) *F. johnsoniae* with daidzein + 50:50 methanol:water 6) *F. johnsoniae* with DMSO + *B. thailandensis* supernatant and 7) *F. johnsoniae* with daidzein + *B. thailandensis* supernatant. A volume of 1.04 µL was added to tubes containing Daidzein (10 mg/ml) or DMSO and a volume of 16.25 µL was added to tubes containing *B. thailandensis* supernatant or 50:50 methanol:water. After additions of solvent or components, 200 µL aliquots from each tube was placed in a 96 well plate (Supplementary Fig. 1). The plate contained 5 independent replicates, 2 technical replicates/independent replicate for conditions 1-3 and 3 tech reps/independent replicate for conditions 4-7 (90 samples). For the remaining 6 wells, 200 µL TSB50 was added to each well as a negative control. An initial absorbance reading (590 nm) was taken on a Tecan Infinite® F500 Multimode Microplate Reader (Tecan Group Ltd., Männedorf, Switzerland). The plate was then incubated for 24 h at 27 °C, 200 rpm. A final absorbance reading (590 nm) was taken after 24 h of incubation. ANOVA was used to compare all the controls and results were insignificant (*p* = 0.544). For this reason, we only used the *F. johnsoniae* culture control (with no added solvents) to serve as a control for a follow-up ANOVA that compared final OD across test conditions. TukeyHSD was performed for post-hoc analysis.

### Experimental evolution

*B. thailandensis* E264 and *F. johnsoniae* UW101 were plated from freezer stocks onto TSA50. Plates were incubated over night at 27 °C. A single, isolated colony of *B. thailandensis* and of *F. johnsoniae* were inoculated as separate cultures in 7 mL of TSB50. Cultures were incubated over night at 27 °C, 200 rpm. The following day, the cultures were pelleted by centrifuged at 5000 rpm for 10 min, the supernatant was removed, and the cultures were resuspended in 1X phosphate buffered saline (PBS). This process was repeated once more. The cultures were resuspended in PBS at a final volume of 5 mL. Ancestral freezer stocks were prepared by adding 750 mL of each culture to 750 mL of 70% glycerol. All freezer stocks were stored at -80 °C. From the remaining cultures, the OD was measured and each culture was diluted to an OD of 0.1 in PBS to prepare the evolution experiments. Ten µL of a culture (OD 0.1) was spotted onto M9 minimal salts agar plates containing 0.2% glucose. Strains were plated both in isolation and co-plated together. When co-plated together, *B. thailandensis* and *F. johnsoniae* were spotted 14 mm apart. Five independent replicates were prepared, resulting in 15 plates total. The plates were wrapped with parafilm and incubated at 27 °C for 1.5 months.

After incubation, we performed a plate passage. Sterilized toothpicks were used to resuspend colonies into 1 mL of PBS. For *B. thailandensis*, we preferentially collected radial colony growth that was growing toward *F. johnsoniae*. A section of radial colony growth was collected for *B. thailandensis* colonies in monocultures. The entirety of the *F. johnsoniae* was removed from each plate. First plate passage freezer stocks were prepared by adding 500 mL of each resuspended culture to 500 mL of 70% glycerol. From the remaining cultures, the OD was measured and each culture was diluted to an OD of 0.1 in PBS. The plating scheme was repeated as previously described while preserving the replicate structure of co-evolving lines (e.g. *B. thailandensis* coevolved replicate 1 was re-plated with *F. johnsoniae* coevolved replicate 1). The plates were wrapped with parafilm and incubated at 27 °C for 1.5 months. This process was repeated for a total of 5 plate passages, resulting in a total experimentation time of 7.5 months.

### Measurements of radial colony growth

Prior to a setting up another plate passage, plates were imaged using a scanner (at 1.5 months). A ruler was placed in the scanner to scale pixels to mm for measurements. Images were uploaded to ImageJ2 for analysis (Rueden et al. 2017; Schneider et al. 2012). Radial growth was determined by measuring the distance from the center of the colony to the furthest point of radial growth. Measurements were uploaded to R for ggplot.

### Whole genome sequencing

All *F. johnsoniae* replicates (monoculture and coevolution experiments) from the 5^th^ plate passage and the *F. johnsoniae* ancestor were plated from freezer stocks onto TSA50 (11 freezer stocks total). These were incubated overnight at 27 °C. The following night, isolated colonies were inoculated into TSB50 medium and incubated over night at 27 °C, 200 rpm. The following morning, DNA was extracted from all 11 cultures using the E.Z.N.A.® Bacterial DNA Kit (Omega Bio-tek, Norcross, GA) according to the manufacturer’s instructions. DNA integrity was assessed from 260/280 and 260/230 ratios using a NanoDrop® ND-1000 UV-Vis Spectrophotometer (Thermo Fisher Scientific, Waltham, MA) and quantified using a qubit 2.0 fluorometer (Invitrogen, Carlsbad, CA, USA). DNA samples were sent to the Microbial Genome Sequencing Center (MiGS, Pittsburgh, PA) for whole genome sequencing. Illumina DNA library preparations were performed at the MiGS facility according to standard operating protocols. Sequencing (2×151bp) was performed on a NextSeq 2000 platform. A minimum of 200 Mbp of sequencing data was obtained with >Q30 reads. Mutations were identified using the breseq (0.33.2) pipeline (Deatherage and Barrick 2014).

### TolC model prediction

The *F. johnsoniae* TolC protein sequence of interest (FJOH_RS06580) was downloaded from NCBI. The protein FASTA file was manually edited to remove 11 amino acids associated with the 33 bp deletion (amino acids 87-97). The protein FASTA file was also manually edited to create the nonsynonymous mutation (G83R). Then all three files were placed into SWISS-MODEL to model the protein structure of TolC (Waterhouse et al. 2018). The template used for rendering all models was SMTL ID : 6wxi.1. The model from TolC wild-type was downloaded and uploaded into Swiss-PdbViewer to highlight amino acids associated with the 33 bp deletion (Guex and Peitsch 1997).

### Construction of mutants in *F. johnsoniae* ancestral strain

Single mutations of interest observed in the coevolved lines were engineered into the ancestral strain. These mutants were constructed following the previously described method (Zhu et al. 2017), with the exception a nested PCR step. All primers used in this study are listed in Supplementary Table 2.

### Nested PCR

DNA extracted from coevolved lines for whole genome sequencing was used as templates for PCR. Coevolved line 1 (JCCE01) was used to amplify the 33 bp deletion in *tolC* (*F. johnsoniae* locus FJOH_RS06580; 261-293 in coding sequence), coevolved line 3 (JCCE03) was used to amplify the nonsynonymous mutation in *tolC* (*F. johnsoniae* locus FJOH_RS06580; G247A in coding sequence), and coevolved line 4 (JCCE04) was used to amplify the base insertion sequence in *ragB*/*susD* (*F. johnsoniae* locus FJOH_RS24865; G at 794 in coding sequence). Nested PCR was performed to obtain enough DNA for restriction digestion and plasmid ligation. For the first round of nested PCR, a 3.3-kbp fragment containing *tolC*Δ33 or *tolC*(G247A) was amplified using primers 1001 and 1002. A 3.5-kbp fragment containing *ragB*794_795insG was amplified using primers 1010 and 1011. The PCR reactions contained reagents and volumes outlined in Supplementary Table 3. PCR conditions were as follows: 98 °C for 30 s, 98 °C for 10 s, 56 °C for 15 s, and 72°C for 70 s, repeated 29 times from step 2, followed by 72°C for 10 min and hold at 4 °C. PCR products were run on a 0.8% agarose gel at 100V for 60 min. PCR bands at the correct fragment sizes were excised from the gel and extracted using Wizard® SV Gel and PCR Clean-Up System (Promega Corporation, Madison WI). PCR products were quantified using a qubit 2.0 fluorometer (Invitrogen, Carlsbad, CA, USA).

For the second round of PCR, a 3.2-kbp fragment containing *tolC*Δ33 or *tolC*(G247A) was amplified using primers 1003 (engineered XbaI site) and 1004 (engineered BamHI site). A 3.1-kbp fragment containing *ragB*794_795insG was amplified using primers 1012 (engineered XbaI site) and 1013 (engineered BamHI site). The PCR reactions contained reagents and volumes outlined in Supplementary Table 4. PCR conditions were as follows: 98 °C for 30 s, 98 °C for 10 s, 56 °C for 15 s, and 72°C for 70 s, repeated 29 times from step 2, followed by 72°C for 10 min and hold at 4 °C. PCR products were run on a 0.8% agarose gel at 100V for 60 min. PCR bands at the correct fragment sizes were excised from the gel and extracted using Wizard® SV Gel and PCR Clean-Up System (Promega Corporation, Madison WI). PCR products were quantified using a qubit 2.0 fluorometer (Invitrogen, Carlsbad, CA, USA).

### Plasmid isolation and purification

A *E. coli* DH5ɑmcr_pYT354 freezer stock was plated on lysogeny broth (LB) agar with ampicillin (100 µg/mL) and incubated overnight at 37 °C. A single colony was inoculated into 5 mL LB with ampicillin (100 µg/mL) and incubated at 37 °C, 200 rpm overnight. Plasmid was extracted and purified the following morning using the E.Z.N.A.® Plasmid DNA Mini Kit I Q-spin (Omega Bio-tek, Norcross, GA) according to the manufacturer’s instructions.

### Restriction enzyme digestion and ligation

Separately prepared restriction enzyme double digestions were performed on purified PCR products from nested PCR round 2 and pYT354. The reaction reagents and volumes are outlined in Supplementary Table 5. Reactions were incubated at 37 °C for 15 min. Reactions were then run on a 0.8% agarose gel at 100V for 60 min. Bands at the correct fragment sizes were excised from the gel and extracted using Wizard® SV Gel and PCR Clean-Up System (Promega Corporation, Madison WI). PCR products were quantified using a qubit 2.0 fluorometer (Invitrogen, Carlsbad, CA, USA).

The PCR fragment containing *tolC*Δ33 was ligated to pYT354 to form pJC101, the PCR fragment containing *tolC* (G247A) was ligated to pYT354 to form pJC102, and the PCR fragment containing *ragB*794_795insG was ligated to pYT354 to form pJC103. Ligation reagents are outlined in Supplementary Table 6. The reactions were incubated at room temperature for 10 min and then heat inactivated for 10 min at 65 °C.

### Preparation of heat shock competent cells

A *E. coli* DH5ɑmcr freezer stock was plated on lysogeny broth (LB) agar and incubated overnight at 37 °C. A single colony was inoculated into 5 mL LB and incubated at 37 °C, 200 rpm overnight. One mL of the overnight cultured was diluted into 100 mL LB and incubated at 37 °C, 200 rpm until the OD reached 0.3 (∼ 2 h). The culture was placed on ice for 15 min. Two 45 mL aliquots were placed into 2, 50 mL falcon tubes and centrifuged at 4 °C, 4000 rpm for 10 min. Supernatant was decanted, and the pellets were resuspended in 45 mL ice-cold 0.1 M CaCl_2._ The resuspended cultures were placed on ice for 30 min. Cultures were then centrifuged at 4 °C, 4000 rpm for 10 min. Supernatant was decanted, and the pellets were resuspended in 4.5 mL ice-cold 0.1 M CaCl_2_ with 15% glycerol. The cultures were then distributed as 50 µL aliquots into microcentrifuge tubes. Competent cells were stored as freezer stocks at -80 °C until ready for use.

### Heat shock transformation

Heat shock competent cells (50 µL) were removed from the freezer and placed on ice for 20 min (4 tubes, 1 for each ligation product and 1 pYT354 plasmid control). Two µL of ligation products (and 1 µL of 10 ng/µL pYT354 plasmid) were added to each tube and placed on ice for an additional 20 min. Cells were then heat shocked for 45 s at 42 °C. Heat shocked cells were placed on ice for 2 min. One mL of super optimal broth (SOC) medium was added to each tube and the tubes were incubated at 37 °C, 200 rpm for 1 h. Cells were pelleted by centrifugation at 4000 g for 2 min at 4 °C. The supernatant was removed (950 µL) and the remaining culture was plated on LB agar containing ampicillin (100 µg/mL). Plates were incubated over night at 37 °C. Successful transformants were inoculated into LB containing ampicillin (100 µg/mL) and incubated overnight. Freezer stocks were made the following morning for triparental conjugation.

### Triparental conjugation and recombinant confirmation

pJC101, pJC103, and pJC104 in recombinant *E. coli* DH5αMCR needed to be transferred to the *F. johnsoniae* ancestral strain. This was introduced into *F. johnsoniae* by triparental conjugation as previously described (Rhodes et al. 2011) using recombinant *E. coli* DH5αMCR, *F. johnsoniae* ancestor, and *E. coli* HB101 (carrying the helper plasmid pRK2013), except that the *sacB* and *ermF*-containing suicide vector was used to select for successful *F. johnsoniae* recombinants (Zhu et al. 2017). *F. johnsoniae* recombinants (JCAC01, JCAC02, JCAC03) were confirmed by sanger sequencing at the Michigan State Genomics Core using primers 1005 and 1006 for *tolC*Δ33 or *tolC*(G247A) recombinants and primers 1014 and 1014 for the *ragB*794_795insG recombinant.

### Acetyl-CoA carboxylase carboxyl transferase subunit alpha (AccA) protein alignment

AccA sequences were download from *F. johnsoniae* UW101 (protein ID: WP_011921560.1) and from *Salmonella enterica* serovar Typhimurium strain LT2 (protein ID: NP_459237.1) on NCBI and concatenated as a text file. The text file was uploaded to T-Coffee (Version_11.00) for protein alignment using default parameters (Di Tommaso et al. 2011). The FASTA alignment file from T-Coffee output was download and used as input for BoxShade (version 3.21) using default parameters. The protein alignment was then uploaded to Inkscape for final edits.

### Re-plating experiments

Strains of interest were plated from freezer stocks onto TSA50. Plates were incubated over night at 27 °C. Single isolated colonies were inoculated as separate cultures in 7 mL TSB50. Cultures were incubated over night at 27 °C, 200 rpm. The following day, the cultures were pelleted by centrifuged at 5000 rpm for 10 min, the supernatant was removed, and the cultures were resuspended in 1X PBS. This process was repeated once more. The cultures were resuspended in PBS at a final volume of 5 mL. The OD was measured and each culture was diluted to an OD of 0.1 in PBS. Ten µL of a culture (OD 0.1) was spotted onto M9 minimal salts agar plates containing 0.2% glucose. Strains were either plated in isolation or co-plated together. When co-plated together, strains were spotted 14 mm apart. Plates were incubated at 27 °C for various amounts of time (see Fig. 6 and Supplementary Figs. 4,5, and 8 legends for details). *B. thailandensis* transposon mutants were acquired from the Manoil lab (Gallagher et al. 2013).

## Acknowledgements

We thank Dr. Mark McBride for sending us the plasmid pYT354 in DH5ɑMCR used for recombinant construction and strain HB101 used for triparental conjugation. We also thank Dr. Yongtao Zhu for his consultation in creating *F. johnsoniae* recombinants. Lastly, we thank members of the Dr. Colin Manoil laboratory for providing transposon mutants.

## Statements and Declarations

### Data and code availability

Supplemental files, sanger sequencing files, and our breseq pipeline are available at [https://github.com/ShadeLab/Paper_Chodkowski_Coevolution_2022]. *F. johnsoniae* whole genome raw sequences files are deposited in the NCBI Sequence Read Archive (BioProject ID PRJNA812898).

### Authors’ contributions

JLC and AS conceived the project. JLC carried out the experiments and data analysis. JLC and AS wrote the manuscript.

## References

Allen, H. K., Donato, J., Wang, H. H., Cloud-Hansen, K. A., Davies, J., & Handelsman, J. (2010). Call of the wild: Antibiotic resistance genes in natural environments. Nature Reviews Microbiology, 8(4), 251–259. https://doi.org/10.1038/nrmicro2312

Amunts, A., Fiedorczuk, K., Truong, T. T., Chandler, J., Peter Greenberg, E., & Ramakrishnan, V. (2015). Bactobolin A binds to a site on the 70S ribosome distinct from previously seen antibiotics. Journal of Molecular Biology, 427(4), 753–755. https://doi.org/10.1016/j.jmb.2014.12.018

Aparna, V., Dineshkumar, K., Mohanalakshmi, N., Velmurugan, D., & Hopper, W. (2014). Identification of natural compound inhibitors for multidrug efflux pumps of Escherichia coli and Pseudomonas aeruginosa using In Silico high-throughput virtual screening and In Vitro validation. PLoS ONE, 9(7), e101840. https://doi.org/10.1371/journal.pone.0101840

Arzanlou, M., Chai, W. C., & Venter, H. (2017). Intrinsic, adaptive and acquired antimicrobial resistance in Gramnegative bacteria. Essays in Biochemistry, 61(1), 49–59. https://doi.org/10.1042/EBC20160063

Baral, B., Akhgari, A., & Metsä-Ketelä, M. (2018). Activation of microbial secondary metabolic pathways: Avenues and challenges. Synthetic and Systems Biotechnology, 3(3), 163–178. https://doi.org/10.1016/j.synbio.2018.09.001

Bavro, V. N., Pietras, Z., Furnham, N., Pérez-Cano, L., Fernández-Recio, J., Pei, X. Y., … Luisi, B. (2008). Assembly and Channel Opening in a Bacterial Drug Efflux Machine. Molecular Cell, 30(1), 114–121. https://doi.org/10.1016/j.molcel.2008.02.015

Baym, M., Lieberman, T. D., Kelsic, E. D., Chait, R., Gross, R., Yelin, I., & Kishony, R. (2016). Spatiotemporal microbial evolution on antibiotic landscapes. Science, 353(6304), 1147–1151. https://doi.org/10.1126/science.aag0822

Blair, J. M. A., Richmond, G. E., & Piddock, L. J. V. (2014). Multidrug efflux pumps in Gram-negative bacteria and their role in antibiotic resistance. http://Dx.Doi.Org/10.2217/Fmb.14.66, 9(10), 1165–1177. https://doi.org/10.2217/FMB.14.66

Bolivar, F., & Backman, K. (1979). Plasmids of Escherichia coli as cloning vectors. Methods in Enzymology, 68(C), 245–267. https://doi.org/10.1016/0076-6879(79)68018-7

Brett, P. J., DeShazer, D., & Woods, D. E. (1998). Burkholderia thailandensis sp. nov., a Burkholderia pseudomallei-like species. International Journal of Systematic Bacteriology, 48(1), 317–320. https://doi.org/10.1099/00207713-48-1-317

Bzymek, M., & Lovett, S. T. (2001). Instability of repetitive DNA sequences: The role of replication in multiple mechanisms. Proceedings of the National Academy of Sciences of the United States of America, 98(15), 8319–8325. https://doi.org/10.1073/pnas.111008398

Castañeda-García, A., Rodríguez-Rojas, A., Guelfo, J. R., & Blázquez, J. (2009). The glycerol-3-phosphate permease GlpT is the only fosfomycin transporter in Pseudomonas aeruginosa. Journal of Bacteriology, 191(22), 6968–6974. https://doi.org/10.1128/JB.00748-09

Colclough, A. L., Scadden, J., & Blair, J. M. A. (2019). TetR-family transcription factors in Gram-negative bacteria: Conservation, variation and implications for efflux-mediated antimicrobial resistance. BMC Genomics, 20(1), 1–12. https://doi.org/10.1186/s12864-019-6075-5

Cuthbertson, L., & Nodwell, J. R. (2013). The TetR Family of Regulators. Microbiology and Molecular Biology Reviews, 77(3), 440–475. https://doi.org/10.1128/mmbr.00018-13

D’Costa, V. M., McGrann, K. M., Hughes, D. W., & Wright, G. D. (2006). Sampling the antibiotic resistome. Science, 311(5759), 374–377. https://doi.org/10.1126/science.1120800

Deatherage, D. E., & Barrick, J. E. (2014). Identification of mutations in laboratory-evolved microbes from nextgeneration sequencing data using breseq. Methods in Molecular Biology, 1151, 165–188. https://doi.org/10.1007/978-1-4939-0554-6_12

Di Tommaso, P., Moretti, S., Xenarios, I., Orobitg, M., Montanyola, A., Chang, J. M., … Notredame, C. (2011). T-Coffee: A web server for the multiple sequence alignment of protein and RNA sequences using structural information and homology extension. Nucleic Acids Research, 39((Web Server issue)), W13–7. https://doi.org/10.1093/nar/gkr245

Du, D., Wang, Z., James, N. R., Voss, J. E., Klimont, E., Ohene-Agyei, T., … Luisi, B. F. (2014). Structure of the AcrAB-TolC multidrug efflux pump. Nature, 509(7501), 512–515. https://doi.org/10.1038/nature13205

Du, D., Wang-Kan, X., Neuberger, A., van Veen, H. W., Pos, K. M., Piddock, L. J. V., & Luisi, B. F. (2018). Multidrug efflux pumps: structure, function and regulation. Nature Reviews Microbiology, 16(9), 523–539. https://doi.org/10.1038/s41579-018-0048-6

Duerkop, B. A., Varga, J., Chandler, J. R., Peterson, S. B., Herman, J. P., Churchill, M. E. A., … Greenberg, E. P. (2009). Quorum-sensing control of antibiotic synthesis in Burkholderia thailandensis. Journal of Bacteriology, 191(12), 3909–3918. https://doi.org/10.1128/JB.00200-09

Ebbensgaard, A. E., Løbner-Olesen, A., & Frimodt-Møller, J. (2020). The role of efflux pumps in the transition from low-level to clinical antibiotic resistance. Antibiotics, 9(12), 1–7. https://doi.org/10.3390/antibiotics9120855

Figurski, D. H., & Helinski, D. R. (1979). Replication of an origin-containing derivative of plasmid RK2 dependent on a plasmid function provided in trans. Proceedings of the National Academy of Sciences of the United States of America, 76(4), 1648–1652. https://doi.org/10.1073/pnas.76.4.1648

Frimodt-Møller, J., & Løbner-Olesen, A. (2019). Efflux-Pump Upregulation: From Tolerance to High-level Antibiotic Resistance? Trends in Microbiology, 27(4), 291–293. https://doi.org/10.1016/j.tim.2019.01.005

Frimodt-Møller, J., Rossi, E., Haagensen, J. A. J., Falcone, M., Molin, S., & Johansen, H. K. (2018). Mutations causing low level antibiotic resistance ensure bacterial survival in antibiotic-treated hosts. Scientific Reports, 8(1), 1–13. https://doi.org/10.1038/s41598-018-30972-y

Gallagher, L. A., Ramage, E., Patrapuvich, R., Weiss, E., Brittnacher, M., & Manoil, C. (2013). Sequence-defined transposon mutant library of Burkholderia thailandensis. MBio, 4(6), e00604–13. https://doi.org/10.1128/mBio.00604-13

Granato, E. T., Meiller-Legrand, T. A., & Foster, K. R. (2019). The Evolution and Ecology of Bacterial Warfare. Current Biology, 29(11), R521–R537. https://doi.org/10.1016/j.cub.2019.04.024

Guex, N., & Peitsch, M. C. (1997). SWISS-MODEL and the Swiss-PdbViewer: An environment for comparative protein modeling. Electrophoresis, 18(15), 2714–2723. https://doi.org/10.1002/elps.1150181505

Hermsen, R., Deris, J. B., & Hwa, T. (2012). On the rapidity of antibiotic resistance evolution facilitated by a concentration gradient. Proceedings of the National Academy of Sciences of the United States of America, 109(27), 10775–10780. https://doi.org/10.1073/pnas.1117716109

Hermsen, R., & Hwa, T. (2010). Sources and sinks: A stochastic model of evolution in heterogeneous environments. Physical Review Letters, 105(24), 248104. https://doi.org/10.1103/PhysRevLett.105.248104

Hibbing, M. E., Fuqua, C., Parsek, M. R., & Peterson, S. B. (2010). Bacterial competition: Surviving and thriving in the microbial jungle. Nature Reviews Microbiology, 8(1), 15–25. https://doi.org/10.1038/nrmicro2259

Kong, D., & Masker, W. (1994). Deletion between direct repeats in T7 DNA stimulated by double-strand breaks. Journal of Bacteriology, 176(19), 5904–5911. https://doi.org/10.1128/jb.176.19.5904-5911.1994

Krishnamoorthy, G., Tikhonova, E. B., Dhamdhere, G., & Zgurskaya, H. I. (2013). On the role of TolC in multidrug efflux: The function and assembly of AcrAB-TolC tolerate significant depletion of intracellular TolC protein. Molecular Microbiology, 87(5), 982–997. https://doi.org/10.1111/mmi.12143

McBride, M. J., Xie, G., Martens, E. C., Lapidus, A., Henrissat, B., Rhodes, R. G., … Stein, J. L. (2009). Novel features of the polysaccharide-digesting gliding bacterium Flavobacterium johnsoniae as revealed by genome sequence analysis. Applied and Environmental Microbiology, 75(21), 6864–6875. https://doi.org/10.1128/AEM.01495-09

Medema, M. H., Kottmann, R., Yilmaz, P., Cummings, M., Biggins, J. B., Blin, K., … Glöckner, F. O. (2015). Minimum Information about a Biosynthetic Gene cluster. Nature Chemical Biology, 11(9), 625–631. https://doi.org/10.1038/nchembio.1890

Meouche, I. E., & Dunlop, M. J. (2018). Heterogeneity in efflux pump expression predisposes antibiotic-resistant cells to mutation. Science, 362(6415), 686–690. https://doi.org/10.1126/SCIENCE.AAR7981Mungan,

M. D., Alanjary, M., Blin, K., Weber, T., Medema, M. H., & Ziemert, N. (2020). ARTS 2.0: Feature updates and expansion of the Antibiotic Resistant Target Seeker for comparative genome mining. Nucleic Acids Research, 48(W1), W546–W552. https://doi.org/10.1093/NAR/GKAA374

Pei, X. Y., Hinchliffe, P., Symmons, M. F., Koronakis, E., Benz, R., Hughes, C., & Koronakis, V. (2011). Structures of sequential open states in a symmetrical opening transition of the TolC exit duct. Proceedings of the National Academy of Sciences of the United States of America, 108(5), 2112–2117. https://doi.org/10.1073/pnas.1012588108

Poole, K. (2004). Efflux-mediated multiresistance in Gram-negative bacteria. Clinical Microbiology and Infection : The Official Publication of the European Society of Clinical Microbiology and Infectious Diseases, 10(1), 12–26. https://doi.org/10.1111/J.1469-0691.2004.00763.X

Rhodes, R. G., Pucker, H. G., & McBride, M. J. (2011). Development and use of a gene deletion strategy for Flavobacterium johnsoniae to identify the redundant gliding motility genes remF, remG, remH, and remI. Journal of Bacteriology, 193(10), 2418–2428. https://doi.org/10.1128/JB.00117-11

Rueden, C. T., Schindelin, J., Hiner, M. C., DeZonia, B. E., Walter, A. E., Arena, E. T., & Eliceiri, K. W. (2017). ImageJ2: ImageJ for the next generation of scientific image data. BMC Bioinformatics, 18(1), 529. https://doi.org/10.1186/s12859-017-1934-z

Rutledge, P. J., & Challis, G. L. (2015). Discovery of microbial natural products by activation of silent biosynthetic gene clusters. Nature Reviews Microbiology, 13(8), 509–523. https://doi.org/10.1038/nrmicro3496

Sandoval-Motta, S., & Aldana, M. (2016). Adaptive resistance to antibiotics in bacteria: A systems biology perspective. Wiley Interdisciplinary Reviews: Systems Biology and Medicine, 8(3), 253–267. https://doi.org/10.1002/wsbm.1335

Schneider, C. A., Rasband, W. S., & Eliceiri, K. W. (2012). NIH Image to ImageJ: 25 years of image analysis. Nature Methods. https://doi.org/10.1038/nmeth.2089

van der Meij, A., Worsley, S. F., Hutchings, M. I., & van Wezel, G. P. (2017). Chemical ecology of antibiotic production by actinomycetes. FEMS Microbiology Reviews, 41(3), 392–416. https://doi.org/10.1093/femsre/fux005

Waglechner, N., McArthur, A. G., & Wright, G. D. (2019). Phylogenetic reconciliation reveals the natural history of glycopeptide antibiotic biosynthesis and resistance. Nature Microbiology, 4(11), 1862–1871. https://doi.org/10.1038/s41564-019-0531-5

Walsh, F. (2013). Investigating antibiotic resistance in non-clinical environments. Frontiers in Microbiology, 4, 19. https://doi.org/10.3389/fmicb.2013.00019

Waterhouse, A., Bertoni, M., Bienert, S., Studer, G., Tauriello, G., Gumienny, R., … Schwede, T. (2018). SWISS-MODEL: Homology modelling of protein structures and complexes. Nucleic Acids Research, 46(W1), W296–W303. https://doi.org/10.1093/nar/gky427

Wozniak, C. E., Lin, Z., Schmidt, E. W., Hughes, K. T., & Liou, T. G. (2018). Thailandamide, a fatty acid synthesis antibiotic that is coexpressed with a resistant target gene. Antimicrobial Agents and Chemotherapy, 62(9), e00463–18. https://doi.org/10.1128/AAC.00463-18

Wu, Y., & Seyedsayamdost, M. R. (2018). The Polyene Natural Product Thailandamide A Inhibits Fatty Acid Biosynthesis in Gram-Positive and Gram-Negative Bacteria. Biochemistry, 57(29), 4247–4251. https://doi.org/10.1021/acs.biochem.8b00678

Yoneyama, H., & Nakae, T. (1993). Mechanism of efficient elimination of protein D2 in outer membrane of imipenem-resistant Pseudomonas aeruginosa. Antimicrobial Agents and Chemotherapy, 37(11), 2385–2390. https://doi.org/10.1128/AAC.37.11.2385

Zhang, Q., Lambert, G., Liao, D., Kim, H., Robin, K., Tung, C. K., … Austin, R. H. (2011). Acceleration of emergence of bacterial antibiotic resistance in connected microenvironments. Science, 333(6050), 1764–1767. https://doi.org/10.1126/science.1208747

Zhu, Y., Thomas, F., Larocque, R., Li, N., Duffieux, D., Cladière, L., … McBride, M. J. (2017). Genetic analyses unravel the crucial role of a horizontally acquired alginate lyase for brown algal biomass degradation by Zobellia galactanivorans. Environmental Microbiology, 19(6), 2164–2181. https://doi.org/10.1111/1462-2920.13699

