## Supplementary for "A coevolution experiment reveals parallel mutations in the AcrA-AcrB-TolC efflux pump that contributes to bacterial antibiotic resistance"

Data and code availability: https://github.com/ShadeLab/Paper_Chodkowski_Coevolution_2022

### Supplementary Tables

**Supplementary Table 1** Summary of *tolC* loci in *F. johnsoniae*

| Locus | Protein ID | Top blastp hit | AA length |
|---|---|---|---|
| FJOH_RS06580 | WP_012023347.1 | TolC family protein | 444 |
| FJOH_RS07030 | WP_012023433.1 | " | 451 |
| FJOH_RS08665 | WP_012023747.1 | " | 461 |
| FJOH_RS14165 | WP_012024794.1 | " | 415 |
| FJOH_RS15250 | WP_012025000.1 | " | 436 |
| FJOH_RS15955 | WP_008463753.1 | " | 415 |
| FJOH_RS16725 | WP_012025212.1 | " | 469 |
| FJOH_RS16800 | WP_012025225.1 | " | 484 |
| FJOH_RS17335 | WP_044047818.1 | " | 426 |
| FJOH_RS20485 | WP_012025935.1 | " | 472 |
| FJOH_RS22150 | WP_044048008.1 | " | 461 |
| FJOH_RS22200 | WP_012026267.1 | " | 412 |
| FJOH_RS22240 | WP_012026275.1 | " | 472 |
| FJOH_RS23175 | WP_012026451.1 | " | 417 |
| FJOH_RS25120 | WP_012026826.1 | " | 479 |
| FJOH_RS25325 | WP_012026867.1 | " | 439 |

**Supplementary Table 2** Primers used in this study

| Primer | Sequence (5’ > 3’) | Description |
|---|---|---|
| 1001 | TTGCTTATTTGGGAGGAACAACA | Used to amplify tolC for nested PCR round 1 |
| 1002 | CATCTGCTTTTGCAGCGATGA | Used to amplify tolC for nested PCR round 1 |
| 1003 | GCTAGTCTAGAGCATCAGTTGAGTTTTCACTGGA | Used for nested PCR round 2 to construct pJC101 and pJC102; XbaI site underlined |
| 1004 | GCTAGGGATCCAAGCTTGCAACCTGGCTTTC | Used for nested PCR round 2 to construct pJC101 and pJC102; BamHI site underlined |
| 1005 | AAATGACGGTCCCATCTCAAA | Used to amplify tolC to confirm successful mutant construction |
| 1006 | CCCATGTAAAACTTCAATGCGT | Used to amplify tolC to confirm successful mutant construction |
| 1010 | TGAGAACCAAAGGCTGGGAA | Used to amplify ragB/susD for nested PCR round 1 |
| 1011 | GGTACATTGTTTTCGGCGCA | Used to amplify ragB/susD for nested PCR round 1 |
| 1012 | GCTAGTCTAGATGGGGATTAACCAGCGACAG | Used for nested PCR round 2 to construct pJC103; XbaI site underlined |
| 1013 | GCTAGGGATCCTTCACCTGCATCGGCAGTTC | Used for nested PCR round 2 to construct pJC103; BamHI site underlined |
| 1014 | ATGCTCCCGCAAAACCAAGA | Used to amplify ragB/susD to confirm successful mutant construction |
| 1015 | ATCAGGACCAGTTGTTGCCG | Used to amplify ragB/susD to confirm successful mutant construction |

**Supplementary Table 3** PCR conditions for nested PCR round 1

| Reagent | Volume (µL) |
|---|---|
| Template (6.25 ng/µL) | 10 |
| Forward/Reverse primers (10 µM) | 2.5 |
| 10 mM dNTPs (Sigma-Aldrich, St. Louis, MO) | 1 |
| Phusion DNA polymerase (New England BioLabs, Ipswich, MA) | 0.5 |
| Phusion 5X buffer (HF buffer for *tolC* and GC buffer for *ragB/susD*) | 9.5 |
| Nuclease-free water | 24 |

**Supplementary Table 4** PCR conditions for nested PCR round 2

| Reagent | Volume (µL) |
|---|---|
| Template (1 ng/µL; PCR product from R1) | 0.5 |
| Forward/Reverse primers (10 µM) | 2.5 |
| 10 mM dNTPs | 1 |
| Phusion DNA polymerase | 0.5 |
| Phusion 5X buffer (HF buffer for *tolC* and GC buffer for *ragB/susD*) | 9.5 |
| Nuclease-free water | 33.5 |

**Supplementary Table 5** Reagents and reaction volumes for restriction enzyme digestion

| Reagent | Volume (µL) |
|---|---|
| Nested PCR R2 products or pYT354 (1 µg/µL) | 1 |
| 10X cutsmart buffer (New England BioLabs, Ipswich, MA) | 5 |
| BamHI-HF (New England BioLabs, Ipswich, MA) | 1 (20 units) |
| XbaI (New England BioLabs, Ipswich, MA) | 1 (20 units) |
| Nuclease-free water | 42 |

**Supplementary Table 6** Reagents and reaction volumes/mass for ligation reactions

| Reagent | Volume/Mass |
|---|---|
| Insert (~3.2 for *tolC*, ~3.1 kbp for *ragB/susD*) | Varied[a] |
| Vector (~7.7 kbp) | 50 ng |
| T4 DNA ligase (New England BioLabs, Ipswich, MA) | 1 µL |
| 10 X T4 DNA ligase buffer (New England BioLabs, Ipswich, MA) | 2 µL |
| Nuclease-free water | Up to 20 µL |

[a]To achieve a 1:3 vector:insert molar ratio, 61.49 ng was used from *tolC*-containing PCR products and 59.37 ng used from *ragB/susD*-containing PCR products.

### Supplementary Figures

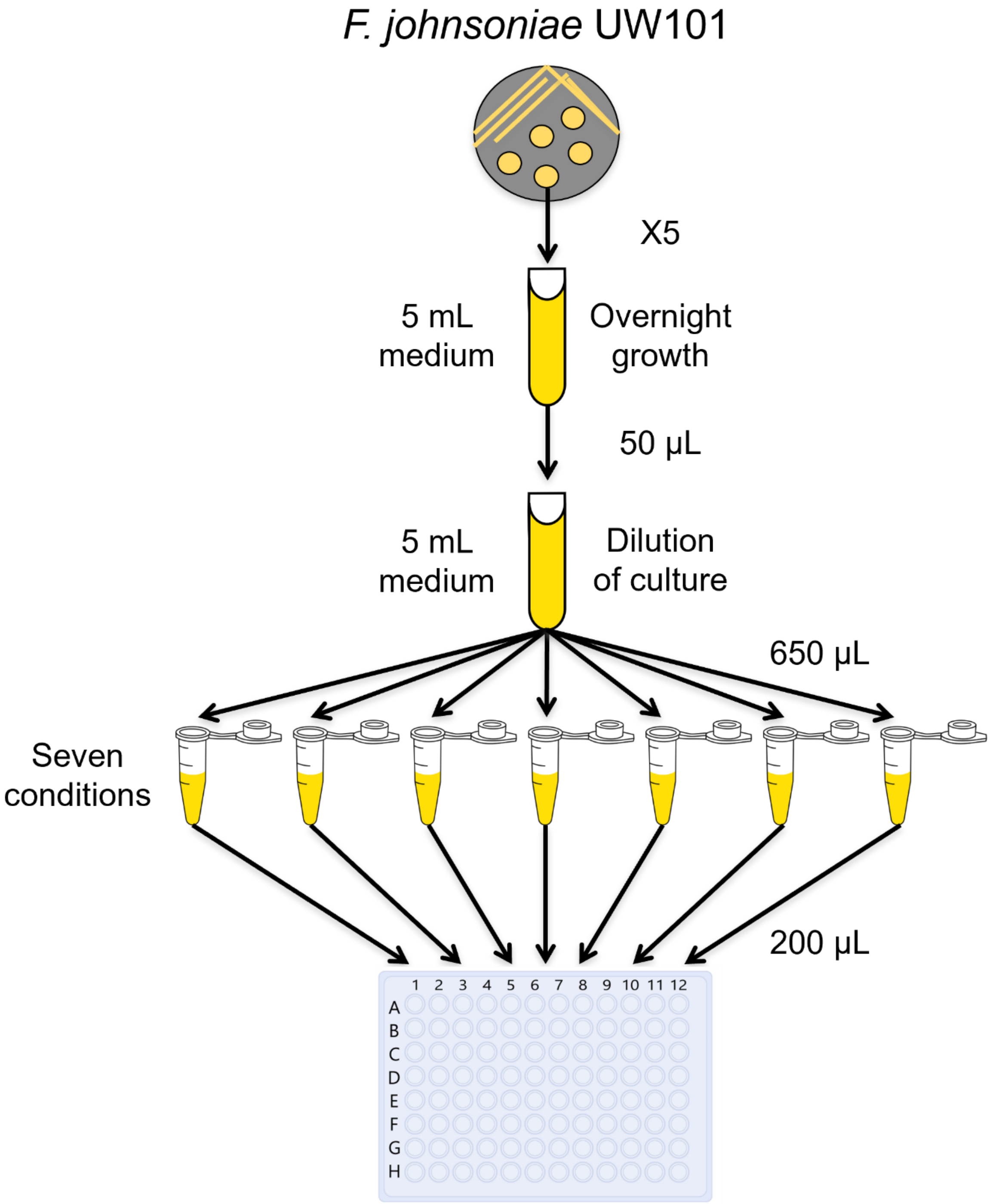


**Supplementary Fig. 1** Schematic of preparation for efflux pump inhibitor experiment.

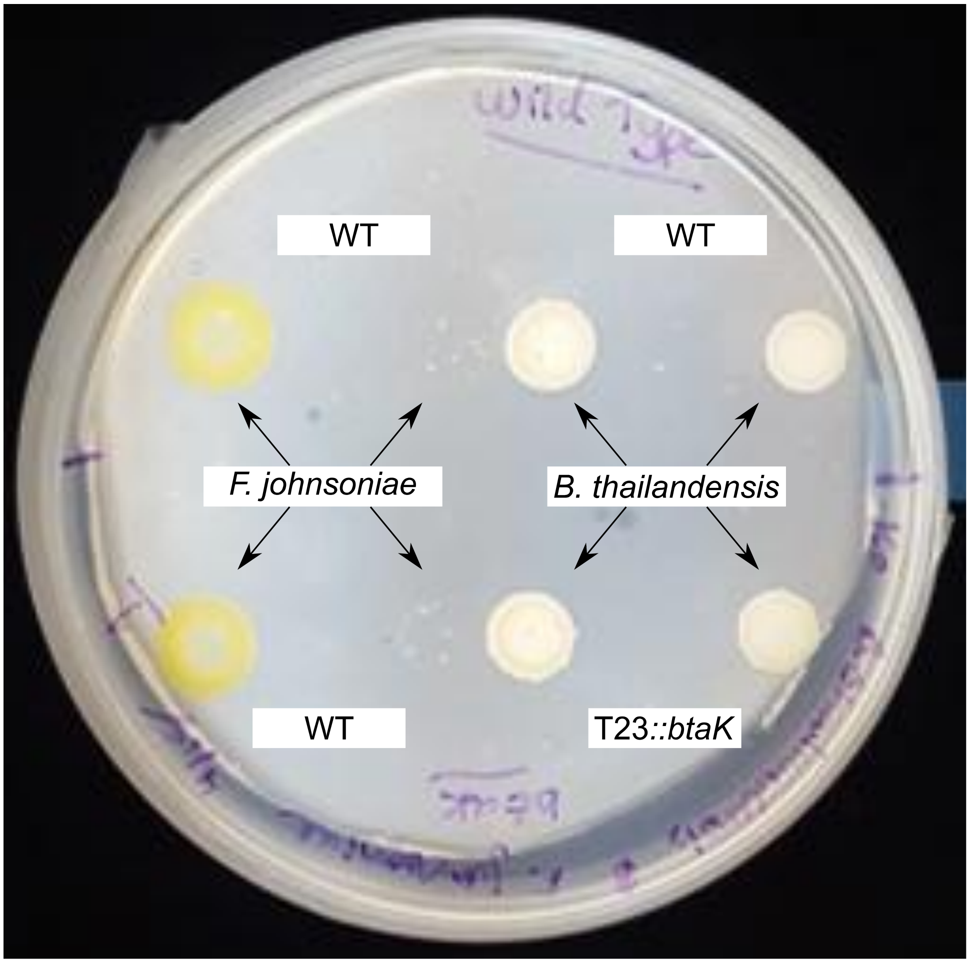


**Supplementary Fig. 2** A *B. thailandensis* btaK::T23 mutant unable to produce bactobolin still inhibits *F. johnsoniae*. *B. thailandensis* WT (top) and *B. thailandensis* btaK::T23 (bottom) was co-plated with *F. johnsoniae* WT. Strains were also plated outside the interspecies interaction zone as controls

Passage *B. thailandensis* *F. johnsoniae*

1

2

3

4

5

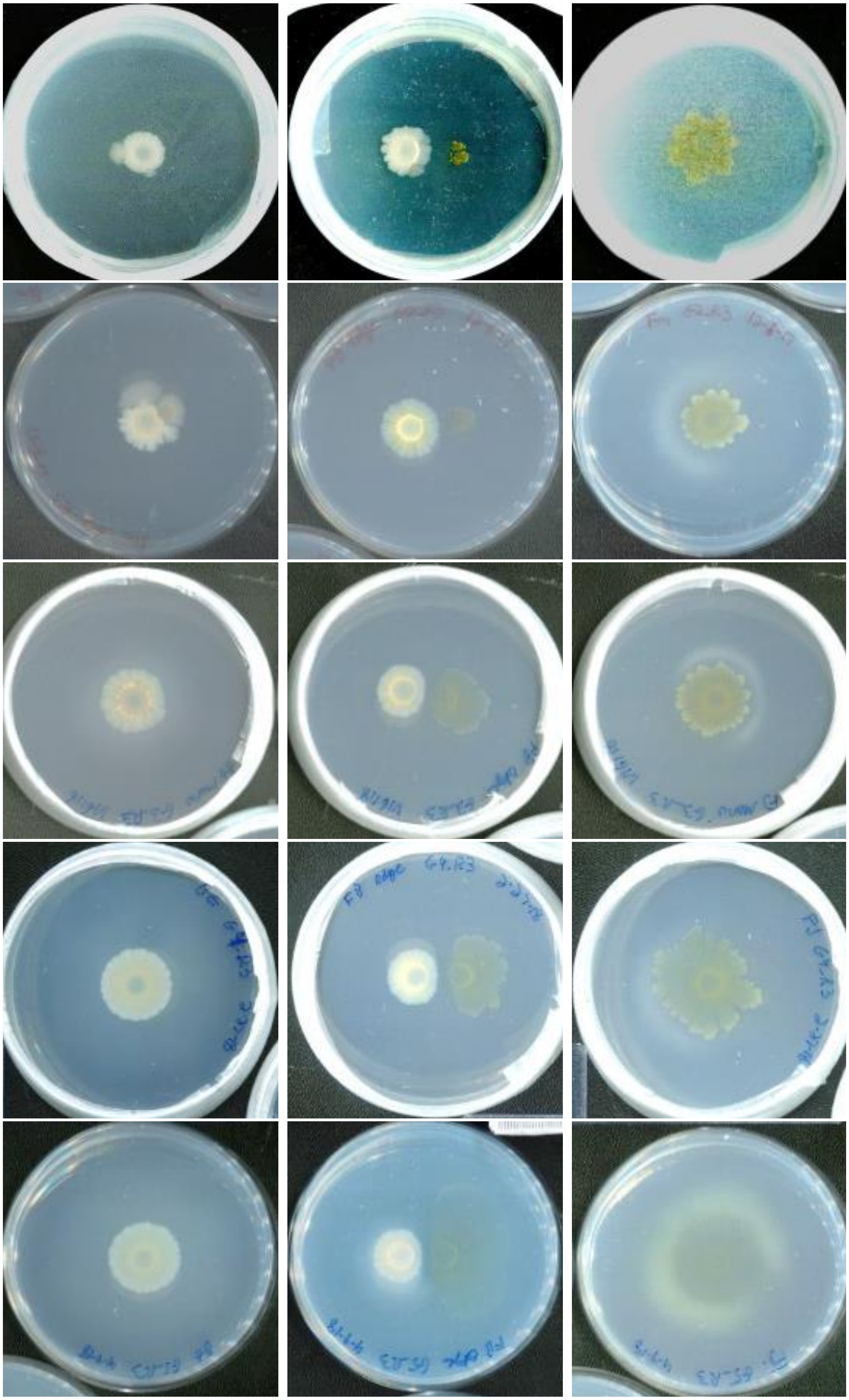

**Supplementary Fig. 3** Colony morphologies and growth success over the (co)evolution experiment. Plate images were taken at 1.5 months after each plate passage. Shown are colony morphologies and growth success of *B. thailandensis* monoculture (column 1), co-plated *B. thailandensis-F. johnsoniae* (column 2), and *F. johnsoniae* monoculture (column 3) for a representative independent replicate (rep 3). Each row is plate passage

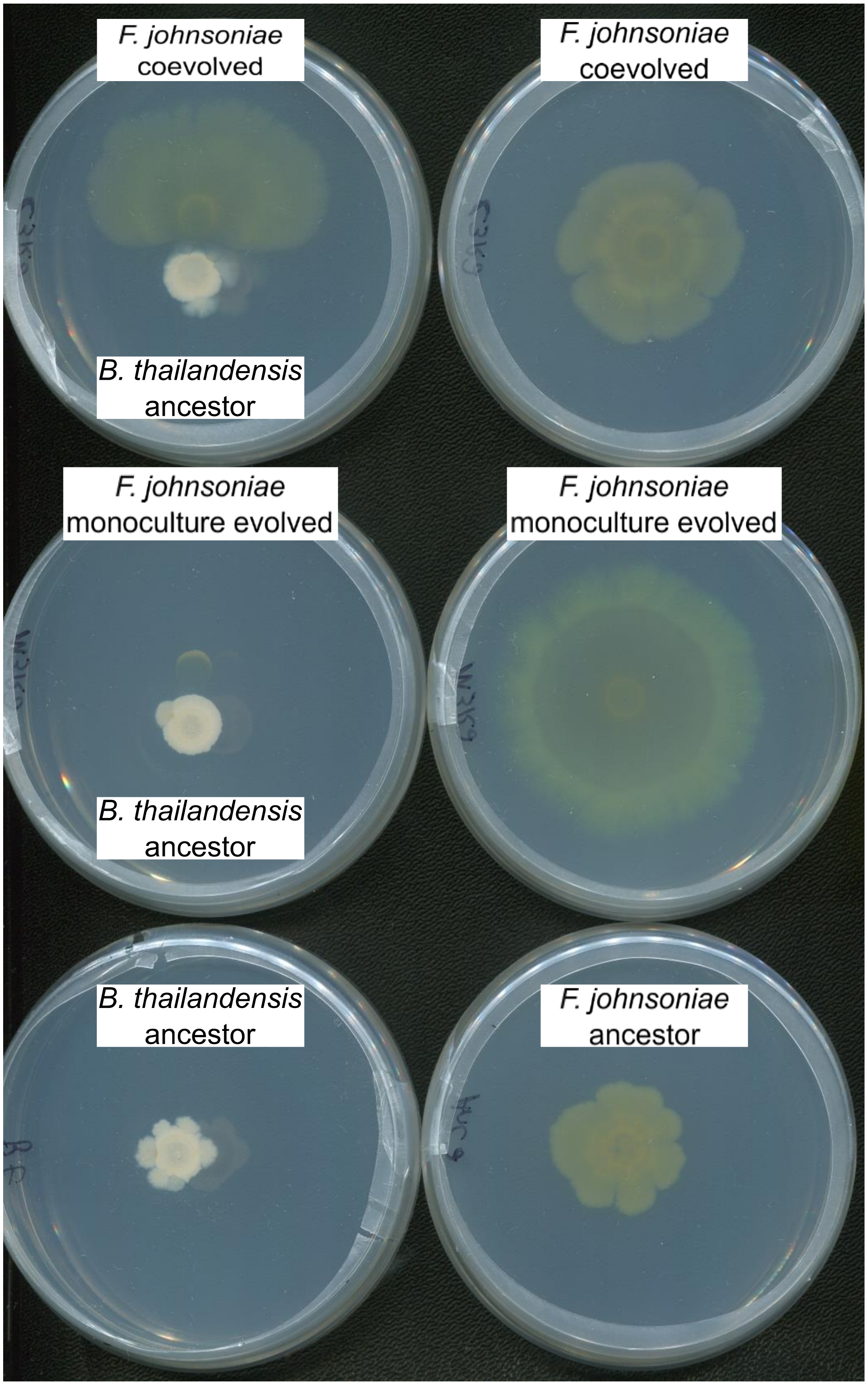
F. johnsoniae coevolved
B. thailandensis ancestor
F. johnsoniae coevolved
F. johnsoniae monoculture evolved
B. thailandensis ancestor
F. johnsoniae monoculture evolved
B. thailandensis ancestor
F. johnsoniae ancestor

**Supplementary Fig. 4** Coevolved *F. johnsoniae* has gained resistance to a *B. thailandensis*-produced antibiotic(s). Coevolved *F. johnsoniae* can grow better in the presence of *B. thailandensis* (column 1, top row) compared to the monoculture evolved *F. johnsoniae* (column 1, middle row). Monocultures are shown as a growth control (column 2, top and middle rows). Shown are evolved lines from one of the independent replicates (rep 3) from the fifth plate passage. Ancestor *F. johnsoniae* and ancestor *B. thailandensis* are shown as additional monoculture controls (bottom row). Images were taken after incubation for 1.5 months

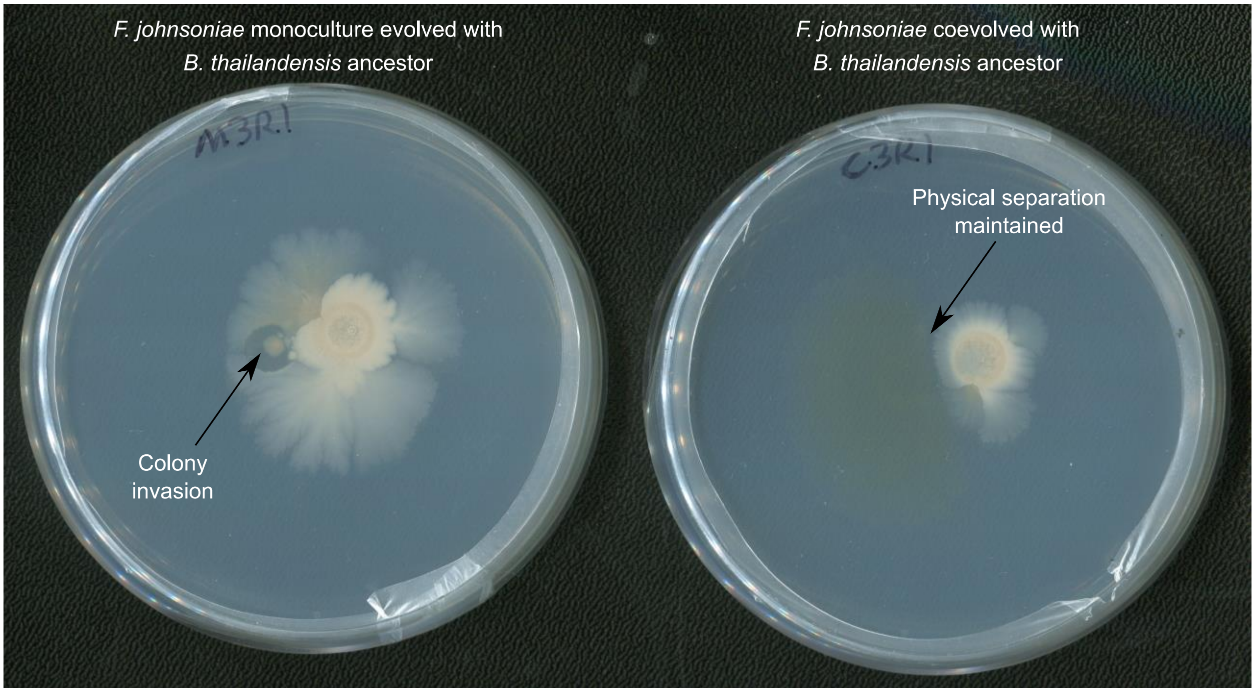


**Supplementary Fig. 5** Coevolved *F. johnsoniae* can resist colony invasion. On each plate, *F. johnsoniae* is on the left and *B. thailandensis* (right) is on the right. The *B. thailandensis* ancestor was co-plated with *F. johnsoniae* evolved monoculture (left plate) and *F. johnsoniae* coevolved (right plate) from the fifth plate passage. Plates were incubated for 2.5 months to allow the chance for physical interactions to occur

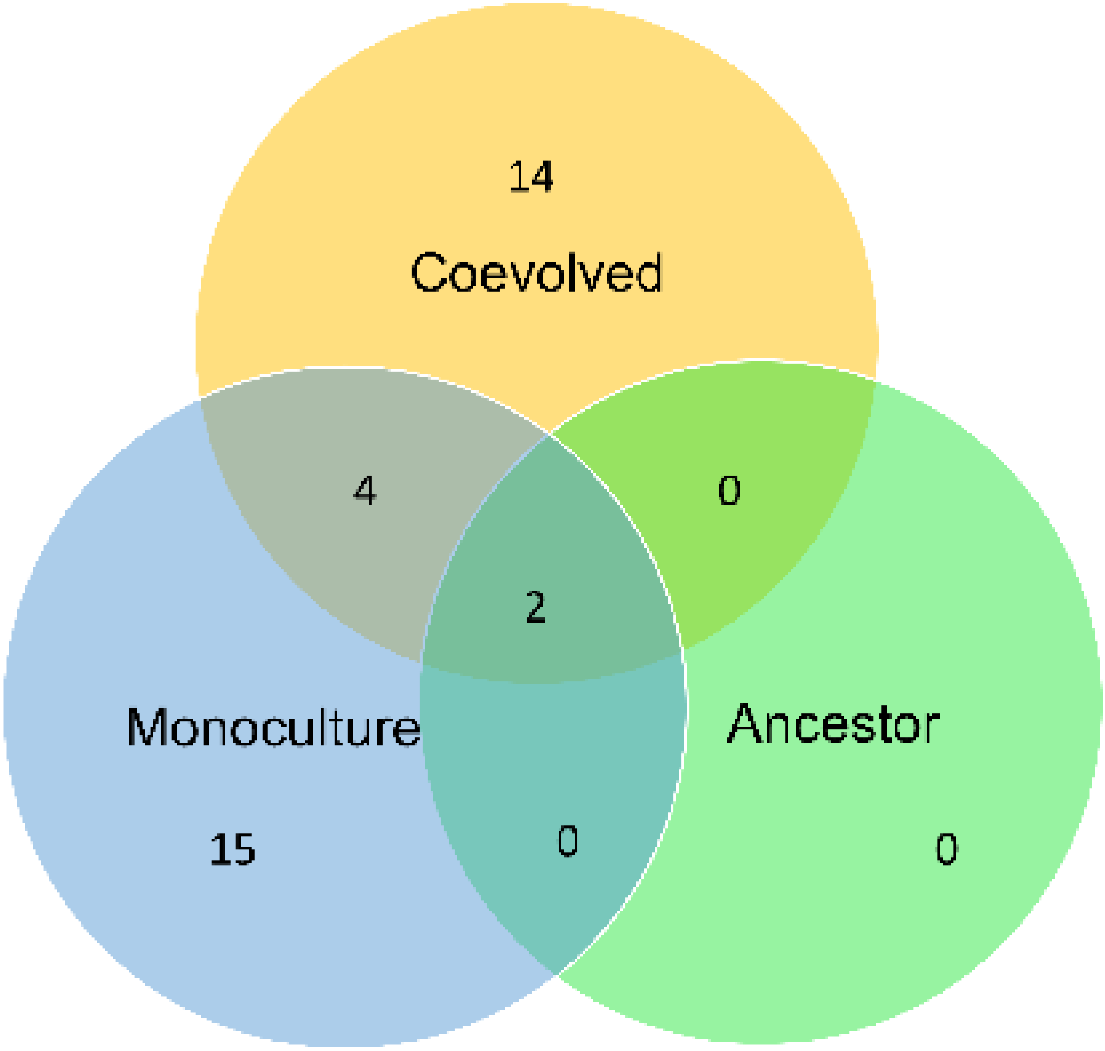


**Supplementary Fig. 6** Coevolved lines acquire unique mutations as a result of interspecies interactions. Shown is a Venn diagram comparing distinctions and overlaps of gene loci where mutations were observed in the ancestor, monoculture evolved lines, and coculture evolved lines

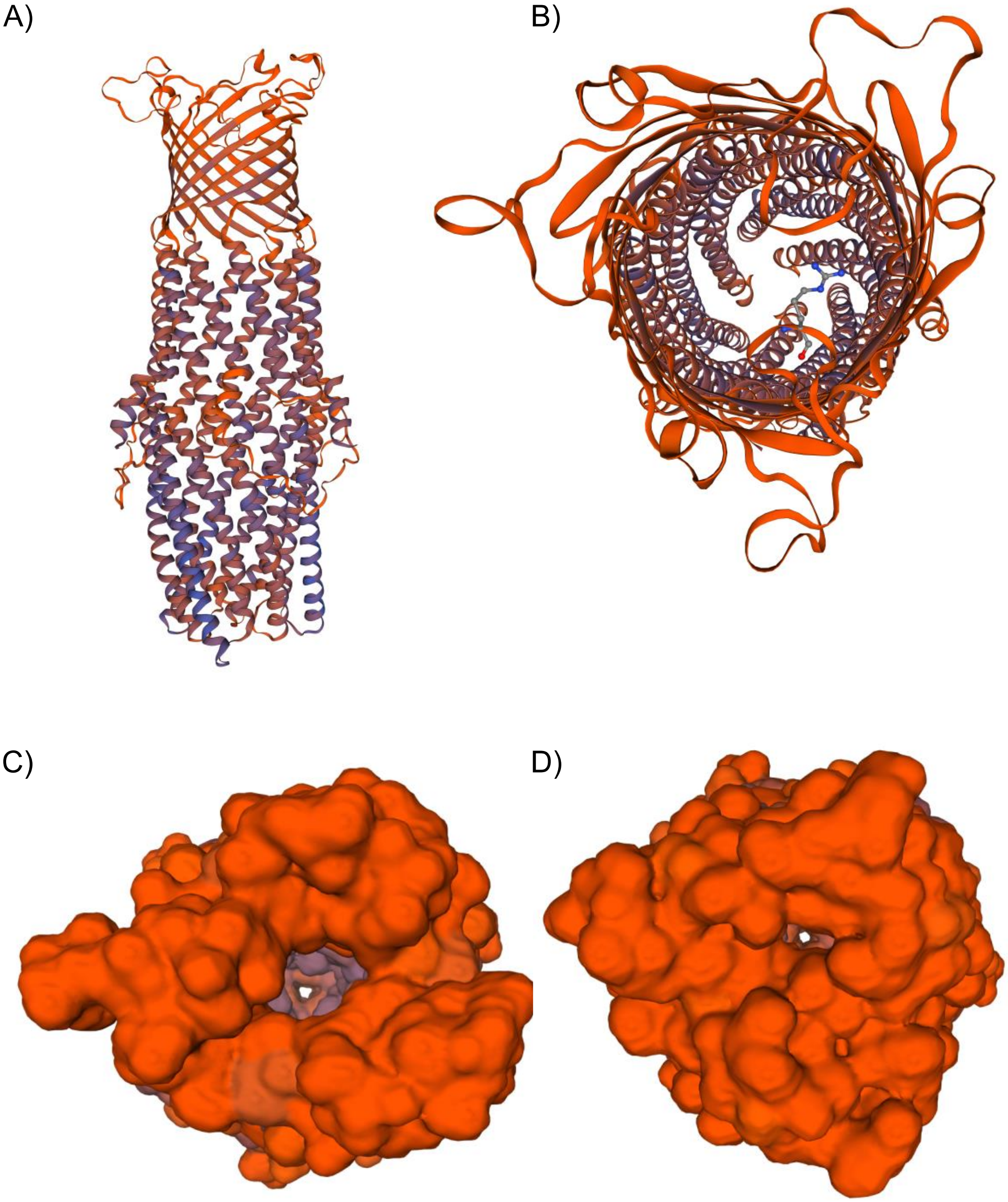


**Supplementary Fig. 7** A nonsynonymous mutation in TolC narrows the opening of the efflux channel. A model of TolC (A) with the G83R nonsynonymous mutation. TolC is rotated +90 about the x-axis in panels C-D such TolC is viewed from top looking down the channel. The G83R residue (B) is located on one of the extracellular loops of TolC. The opening of the efflux channel in WT TolC (C) is predicted to narrow due to G83R (D)

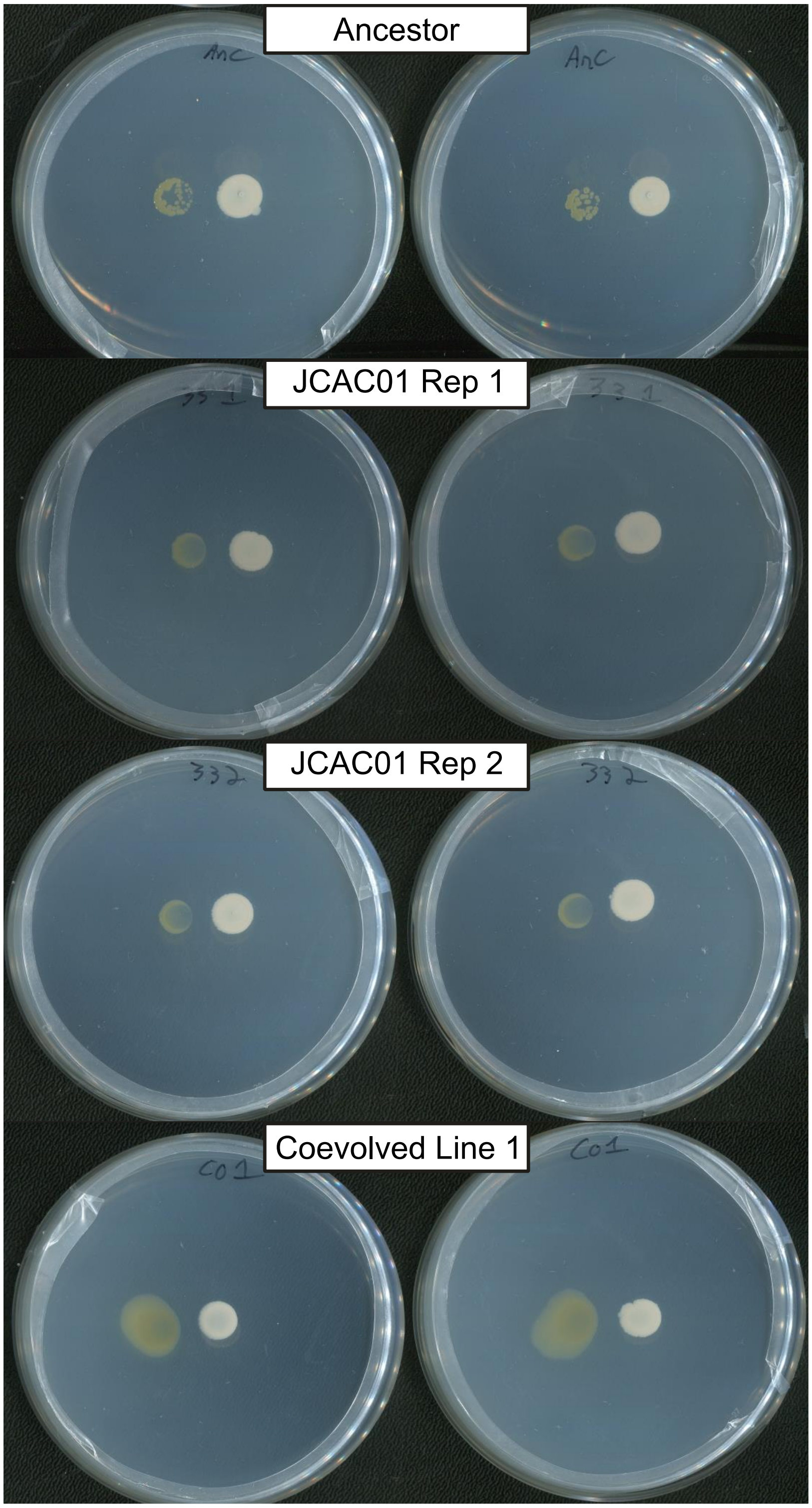


**Supplementary Fig. 8** *F. johnsoniae* recombinants display antibiotic resistance, but not to the extent observed in coevolved lines. The 33 bp deletion in FJOH_RS06580 was placed into the *F. johnsoniae* ancestor and co-plated with *B. thailandensis*. Two confirmed successful recombinants (JCAC01, replicates 1&2) are less inhibited by *B.*

*thailandensis* compared to the *F. johnsoniae* ancestor but are more inhibited compared to the coevolved line from which FJOH_RS06580 was amplified to create the recombinants. All strains were co-plated with the *B. thailandensis* ancestor. Plates were imaged after a month of incubation
